# *Burkholderia* from fungus gardens of fungus-growing ants produce antifungals that inhibit the specialized parasite *Escovopsis*

**DOI:** 10.1101/2021.01.22.427492

**Authors:** Charlotte B. Francoeur, Daniel S. May, Margaret W. Thairu, Don Q. Hoang, Olivia Panthofer, Tim S. Bugni, Mônica T. Pupo, Jon Clardy, Adrián A. Pinto-Tomás, Cameron R. Currie

## Abstract

Within animal associated microbiomes, the functional roles of specific microbial taxa are often uncharacterized. Here, we use the fungus-growing ant system, a model for microbial symbiosis, to determine the potential defensive roles of key bacterial taxa present in the ants’ fungus gardens. Fungus gardens serve as an external digestive system for the ants, with mutualistic fungi in the genus *Leucoagaricus* spp. converting plant substrate into energy for the ants. The fungus garden is host to specialized parasitic fungi in the genus *Escovopsis*. Here, we examine the potential role of *Burkholderia* spp. that occur within ant fungus gardens in inhibiting *Escovopsis.* We isolated members of the bacterial genera *Burkholderia* spp. and *Paraburkholderia* spp. from 50% of the 52 colonies sampled, indicating that the family *Burkholderiaceae* are common fungus garden inhabitants of a diverse range of fungus-growing ant genera. Using antimicrobial inhibition bioassays, we found that 28 out of 32 isolates inhibited at least one *Escovopsis* strain with a zone of inhibition greater than 1 cm. Genomic assessment of *Burkholderiaceae* isolates indicated that isolates with strong inhibition all belonged to the genus *Burkholderia* and contained biosynthetic gene clusters that encoded the production of two antifungals: burkholdine1213 and pyrrolnitrin. Organic extracts of cultured isolates confirmed these compounds as responsible for antifungal activity that inhibit *Escovopsis* but, at low concentrations, not *Leucoagaricus* spp. Overall, these new findings, combined with previous evidence, suggest that members of the fungus garden microbiome play an important role in maintaining the health and function of the fungus-farming ant colony.

**IMPORTANCE:** Many organisms partner with microbes to defend themselves against parasites and pathogens. Fungus-growing ants must protect *Leucoagaricus* spp., the fungal mutualist that provides sustenance for the ants, from a specialized fungal parasite, *Escovopsis* spp. The ants take multiple approaches, including weeding their fungus gardens to remove *Escovopsis* spores, as well as harboring *Pseudonocardia* that produce antifungals that inhibit *Escovopsis.* In addition, a genus of bacteria commonly found in fungus gardens, *Burkholderia* spp., is known to produce secondary metabolites that inhibit *Escovopsis* spp. In this study, we isolated *Burkholderia* spp. from fungus-growing ants, assessed the isolates’ ability to inhibit *Escovopsis* spp., and identified two compounds responsible for inhibition. Our findings suggest that *Burkholderia* spp. are often found in fungus gardens, adding another possible mechanism within the fungus-growing ant system to suppress the growth of the specialized parasite *Escovopsis*.

## INTRODUCTION

Symbiotic associations are ubiquitous. Organisms do not live in isolation, rather, they exist in complex communities consisting of variable macro- and micro-organisms. These symbiotic interactions are known to play a fundamental role in shaping life on earth and can range from transient to obligate and parasitic to beneficial associations (1). Microbial symbioses span this range and fill a variety of important roles within hosts, with substantial work done specifically in association with insect hosts. Research on beneficial microbes in insects has typically focused on the role of symbionts in providing nutrients to their host (2, 3). However, in recent years it has become clear that microbes often play a critical role in mediating interactions between insects and pathogens or parasites (4, 5). Microbes can provide defense through competitively excluding pathogenic microbes, priming the host immune system, and by producing compounds that protect the host (6–8). In this study, we use the fungus-growing ant system to explore a bacterially mediated antifungal defensive symbiosis.

Fungus-growing ants are a well-studied example of a multi-partite symbiosis (Figure 1). Fungus-growing ants (Hymenoptera: Formicidae: Attina) thrive in the Neotropics and consist of 20 genera and approximately 250 species (9, 10) that have formed ancient and highly evolved mutual symbioses with both fungi and bacteria (11). The ants cultivate fungi in the genus *Leucoagaricus* spp. (Basidiomycota: Agaricales: Agaricaceae) as a food source. Fungus-growing ants bring organic substrate to structures known as fungus gardens (Figure 1B), where *Leucoagaricus* spp. principally degrades the substrate and produces usable energy for the ants (12, 13). In addition, a consistent bacterial community composed primarily of Proteobacteria exists within fungus gardens (14–18). In leaf-cutter ants, the bacterial community has been shown to help degrade plant secondary compounds and aid in nitrogen acquisition for the ant through biological nitrogen fixation (19, 20). Though the functions of some of the bacterial taxa have been described, the roles of many bacterial fungus garden members are unknown.

**Figure 1.**
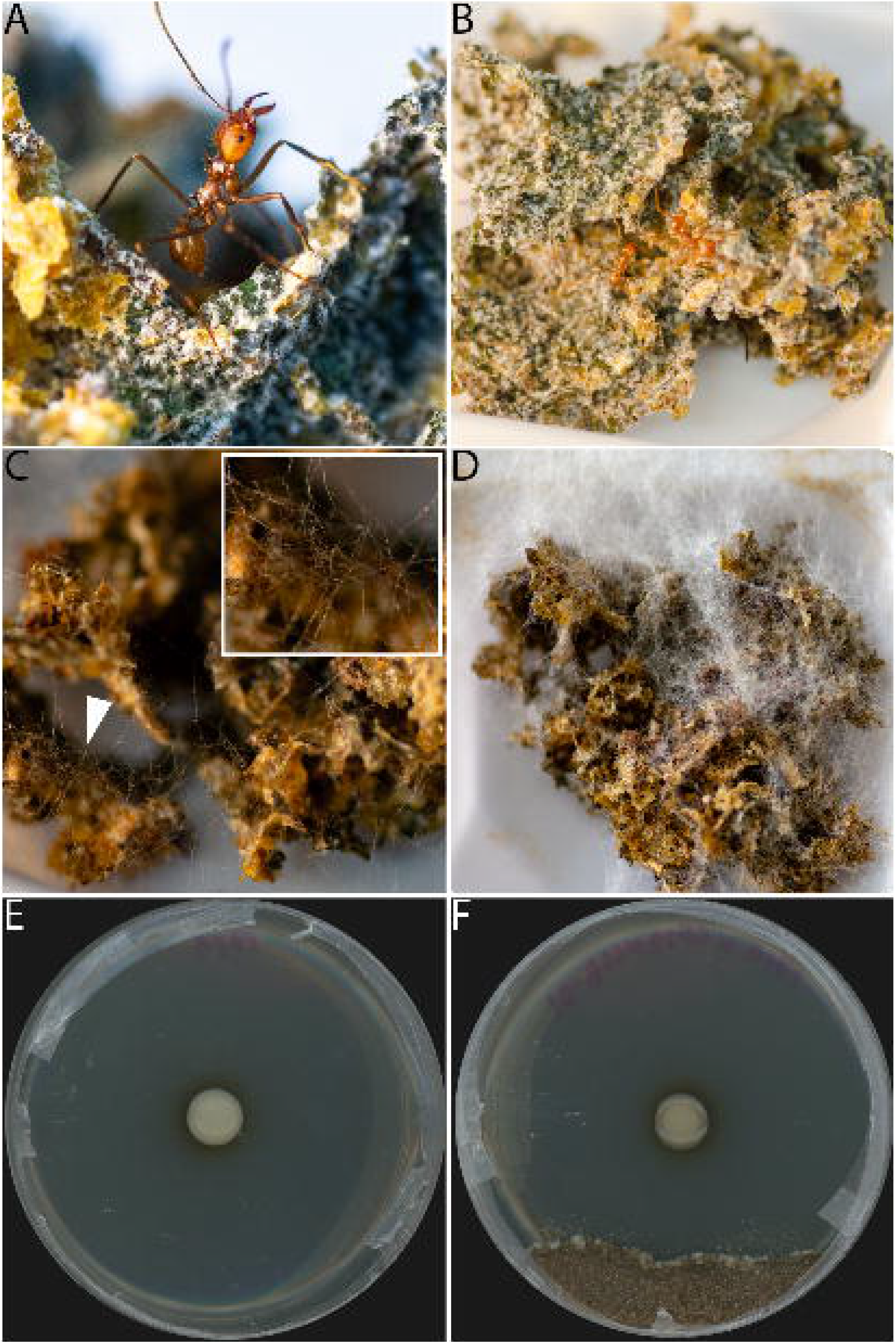
Infection of an *Atta cephalotes* colony by *Escovopsis* sp. CF180408-01 and *in vitro* Petri plate antimicrobial inhibition bioassay with ICBG1719. Healthy fungus garden with no *Escovopsis* infection (A-B). Day 3 post-infection of fungus garden with *Escovopsis* (C). Day 7 post-infection of fungus garden with *Escovopsis* (D). Petri plate with only *Burkholderia* sp. ICBG1719 growing (E) and Petri plate inhibition bioassay of ICBG1719 against *Escovopsis* sp. CF180408-01 with clear zone of inhibition. Pictures A-D taken by Caitlin Carlson.

The fungus garden is threatened by a specialized pathogen, the parasitic fungus, *Escovopsis* (Ascomycota: Hypocreales: Hypocreaceae) (21). In many lineages of fungus-growing ants, the growth of *Escovopsis* is inhibited by Actinobacteria in the genus *Pseudonocardia*, defensive symbionts found on the exoskeleton or occurring within specialized structures on the ants (22–24). However, in the two fungus-growing ant genera *Atta* spp. and *Sericomyrmex* spp. this defensive symbiosis with *Pseudonocardia* has been secondarily lost (23, 24). Despite the lack of the symbiont *Pseudonocardia*, the fungus gardens are not overrun with *Escovopsis*, suggesting other ways of controlling *Escovopsis*. The ant behaviors of weeding and grooming are known to be crucial for suppressing *Escovopsis* (25, 26). In addition, there is evidence that antifungal-producing bacteria may colonize the fungus garden and provide some level of inhibition against *Escovopsis* (27). Santos and colleagues isolated *Burkholderia* consistently from *Atta sexdens* fungus gardens (22/57 colonies) and identified isolates that could inhibit the growth of *Escovopsis*, suggesting a potential role for garden bacteria in defense of fungus gardens.

The fungus gardens’ Proteobacteria-dominant bacterial community is known to include members of the family *Burkholderiaceae* (14–18, 27). *Burkholderiaceae* have diverse metabolic capabilities and inhabit a broad range of ecological niches (28). Two closely-related genera within *Burkholderiaceae*, *Burkholderia* spp. and *Paraburkholderia* spp., have been found and characterized in the context of mammalian and plant pathogenesis, nitrogen fixation, bioremediation, plant growth stimulation, and/or in close association with fungi and insects. These symbiotic *Burkholderia* and *Paraburkholderia* can also produce secondary metabolites that are important in ecological interactions. For example, *Paraburkholderia rhizoxinica* reside in the hyphae of the fungal plant pathogen, *Rhizopus microsporus*. *P. rhizoxinica* produces the antimitotic macrolide that is converted into rhizoxin, which is the causative agent of rice seedling blight (29, 30). Lagriinae beetles depend on *Burkholderia* symbionts for the protection of their eggs. Specifically, *Burkholderia gladioli* produces a blend of antibiotics, including toxoflavin, caryoynencin, lagriene, and sinapigladioside, which protect the egg stage of the beetles against pathogenic microbes (31, 32). Additionally, *Burkholderia* isolates that produce pyrrolnitrin, a characterized antifungal, have been used as biocontrol agents for plant fungal pathogens (33). Finally, as mentioned previously, the study by Santos and colleagues (2004) suggests that antifungal producing *Burkholderia* may play a role in suppressing the fungus garden parasite *Escovopsis* in *Atta sexdens*.

Here, we conduct the first comprehensive investigation of the functional role of fungus garden associated *Burkholderiaceae* isolates obtained from colonies of fungus-growing ants that span major clades in the basal and derived lineages, to test the hypothesis that fungus garden associated bacteria can provide protection against the system’s specialized parasite, *Escovopsis*. First, we sampled 52 fungus-growing ant fungus gardens that span eight different ant genera to isolate *Burkholderiaceae*. We then tested the ability of a subset of *Burkholderiaceae* isolates to inhibit a panel of 11 fungi, including eight strains of the specialized parasite *Escovopsis* that were isolated from five different genera of fungus-growing ants. To identify potential biosynthetic gene clusters involved in *Escovopsis* inhibition, we sequenced the genomes of 30 *Burkholderiaceae* isolates and performed antiSMASH and BiG-SCAPE analyses. Then, to confirm production of antifungals, organic extracts were prepared from a subset of isolates and analytical chemistry techniques were used to identify antifungals. Finally, the organic extracts were also used to assess the inhibition against *Escovopsis* spp. and six strains of *Leucoagaricus* spp.

## RESULTS

### *Burkholderiaceae* are consistently present in the fungal gardens of different lineages of attine ants

*Burkholderiaceae* were frequently isolated from fungus gardens of fungus-growing ants, indicating that they are common residents of fungus gardens. Nutrient-rich, non-selective media was used for the Brazilian fungus garden bacteria isolations and 35% of colonies contained at least one *Burkholderiaceae* isolate (Table 1). *Burkholderiaceae* selective media was used for Costa Rican fungus garden bacteria isolations and 71% of colonies contained at least one *Burkholderiaceae* isolate (Table 1).

**Table 1.** Summary of ant colony collections by geographic location. The number of colonies that had a *Burkholderiaceae* isolate is indicated in parentheses. * indicate two genera, *Atta* and *Sericomyrmex*, that have secondarily lost *Pseudonocardia*.

| Ant Genus | Brazil | Costa Rica |
| --- | --- | --- |
| *Atta* * | 10 (2) | 11 (7) |
| *Acromyrmex* | 3 (0) | 1 (0) |
| *Paratrachymyrmex* | 5 (3) | 2 (2) |
| *Mycetomoellerius* | 0 | 2 (1) |
| *Sericomyrmex* * | 0 | 5 (5) |
| *Mycetophylax* | 2 (2) | 0 |
| *Cyphomyrmex* | 1 (0) | 0 |
| *Mymicocrypta* | 1 (0) | 0 |
| *Apterostigma* | 5 (3) | 0 |
| Unidentified Attini | 4 (1) | 0 |
| Total Colonies | 31 (11) | 21 (15) |

In total, 86 isolates were obtained from these 52 ant colonies and whole 16S rRNA gene sequences aligned with *Burkholderia* spp. as the top BLAST hit (Table S1, Dataset S1) for all isolates. A subset of 30 isolates were selected for whole genome sequencing that represented unique species-level 16S rRNA gene BLAST hits, such that if two isolates from the same fungus garden sample matched the same species, one was randomly selected for sequencing (Table S2). Phylogenetic and Average Nucleotide Identity (ANI) analyses indicated that isolates grouped with both *Burkholderia* spp. and *Paraburkholderia* spp. isolates (Figure S1, Dataset S2), contrary to the original BLAST search with the 16S rRNA gene. Both 16S rRNA gene and whole genome phylogenies (Figure S1) indicate that the bacterial isolates from fungus gardens fall among multiple clades, including different lineages within *Burkholderia* spp., such as plant pathogens (*B. gladioli*), the *B. cepacia* complex, nitrogen-fixing and plant-associated (*B. mimosarum, B. nodosa, B. xenovorans B. phytofirmans)*, as well as plant-, rhizosphere-, and soil-associated *Paraburkholderia* species. ANI analysis confirmed the variety of *Burkholderiaceae* in fungus gardens. 18 out of 30 sequenced isolates shared ≥ 95% ANI with *B. gladioli* (5/18), *B. lata* (9/18), *B. ambifaria* (1/18)*, B. cepacia* (1/18)*, B. seminalis* (1/18), or *Paraburkholderia tropica* (1/18), indicating a range of different characterized *Burkholderiaceae* species. The other 12 isolates shared between 89% and 94% ANI with other *Burkholderiaceae* isolates, such as *B. ubonensis, B. pyrroccinia, P. eburnea,* and *P. caribensis* (Table S2, Dataset S2).

### Multiple fungus garden *Burkholderiaceae* isolates inhibit *Escovopsis*

In order to assess the potential of fungus garden associated *Burkholderiaceae* to inhibit *Escovopsis*, an *in vitro* Petri plate antimicrobial inhibition bioassay with 19 genome-sequenced *Burkholderiaceae* isolates from Brazil and Costa Rica was conducted against a panel of 11 fungi: eight *Escovopsis* isolates, *Aspergillus flavus* (Ascomycota: Eurotiales: Trichocomaceae)*, Fusarium oxysporum* (Ascomycota: Hypocreales: Nectriaceae), and *Trichoderma* sp. (Ascomycota: Hypocreales: Hypocreaceae). After 13 days of co-incubation, all 19 *Burkholderiaceae* isolates had at least one zone of inhibition (ZOI) greater than 0 cm against at least one *Escovopsis* strain (Figure 2, Figure S2-S3). Another set of *in vitro* antimicrobial inhibition bioassays with 13 additional isolates from *Sericomyrmex* colonies in Costa Rica were conducted against a panel of 6 *Escovopsis* strains and the same three non-*Escovopsis* fungi noted above. Similar results were found with the 13 additional isolates; all 13 isolates had at least one ZOI greater than 0 cm against at least one *Escovopsis* strain (Figure S3). *Burkholderia* isolates SID20373, ICBG1719, ICBG1720, ICBG1724, and ICBG1735 inhibited all *Escovopsis* strains significantly more (P<0.001, average zone of inhibition ≥ 2.45 cm; Figure 2), than the other 27 *Burkholderiaceae* isolates surveyed (Figure 2, Figure S3, Table S3). For clarity, the five isolates listed above that inhibited *Escovopsis* strongly will be referred to as strong inhibitory isolates. *Aspergillus flavus, T.* sp., and *F. oxysporum*, were not inhibited by any of the *Burkholderiaceae* isolates.

**Figure 2.**
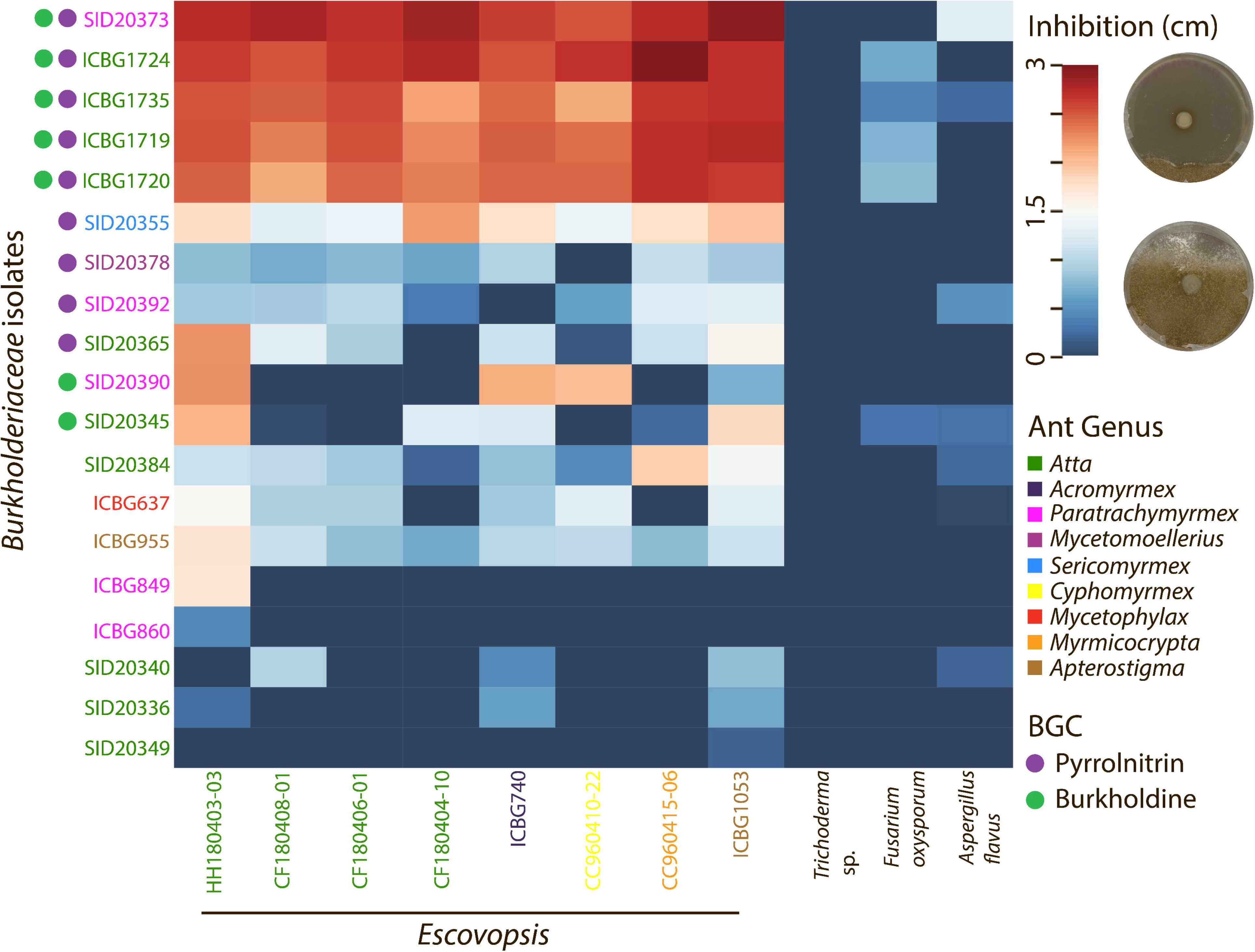
*Burkholderiaceae-Escovopsis* Petri plate antimicrobial inhibition bioassays indicated variable levels of inhibition of *Escovopsis* spp. Zones of inhibition (ZOI) were measured 13 days after inoculation with *Escovopsis* spp. *Burkholderiaceae* and *Escovopsis* isolates are color coded by the ant colony they were isolated from. Circles to the left of the *Burkholderiaceae* isolates indicate presence of pyrrolnitrin (purple) or burkholdine (green) biosynthetic gene clusters.

**Figure 3.**
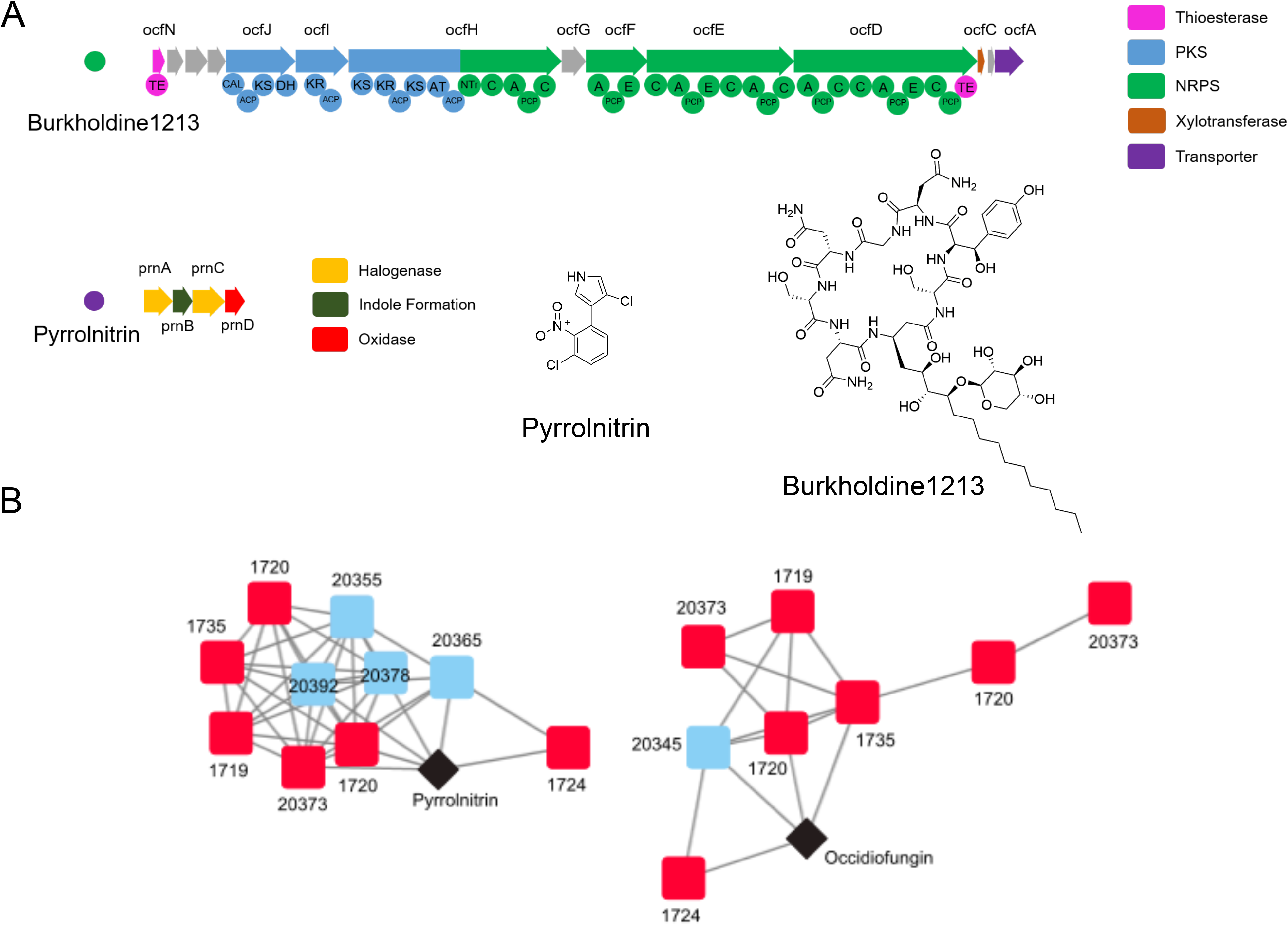
Identification of pyrrolnitrin and burkholdine1213 biosynthetic gene clusters in inhibitory isolates. (A) genetic architecture of biosynthetic gene clusters encoding the production of burkholdine1213 and pyrrolnitrin identified by AntiSMASH 4.0. Domains of the PKS/NRPS hybrid biosynthetic gene cluster are shown beneath the gene representations for burkholdine1213. (B) BiGSCAPE network analysis of the two biosynthetic gene clusters present in *Burkholderia* isolates. Red squares are strongly inhibitory *Burkholderia* isolates, blue squares are less inhibitory isolates, and black diamonds are database matches to the MIBiG biosynthetic gene cluster database.

### Strong inhibitory *Burkholderia* isolates are predicted to have two antifungal biosynthetic gene clusters

To identify secondary metabolites that might be responsible for the inhibitory activity, the sequenced genomes of each *Burkholderiaceae* isolate were submitted to antiSMASH v4.0 for the detection of biosynthetic gene clusters (BGCs), groupings of genes that encode the production of a secondary metabolite. BiGSCAPE was used to compare the presence and absence of BGCs across isolates and to search for correlations between the presence of BGCs and *Escovopsis* inhibition. Two BGCs were identified in the genomes of all five of the inhibitory *Burkholderia* isolates. These BGCs had high similarity by BLASTP to characterized BGCs in the MIBiG database: the BGC for pyrrolnitrin and the BGC for occidiofungin. Pyrrolnitrin is an antifungal alkaloid biosynthesized from tryptophan and initially isolated from *Pseudomonas* spp. (34). Occidiofungin is a hybrid nonribosomal peptide/polyketide antifungal glycopeptide and an analog of the burkholdines (35, 36) (Figure 3). *Burkholderia* spp. isolates that did not strongly inhibit *Escovopsis* contained BGCs for only pyrrolnitrin (SID20355, SID20378, SID20392, SID20365) or only antifungal glycopeptides (SID20390, SID20345) or neither of these BGCs. None of the *Paraburkholderia* spp. isolates (SID20336 and others) demonstrated inhibition against *Escovopsis* and were not predicted to contain either antifungal BGC. The distribution of the two biosynthetic gene clusters across *Burkholderia* isolates is demonstrated in Figure 2.

### *Burkholderia* extracts containing pyrrolnitrin and burkholdine1213 replicate Petri plate inhibition assay

To determine if the antifungal compounds were being produced by the strongly inhibitory isolates, we made organic extracts from *Burkholderia* isolates predicted to have both of the BGCs that encode the production of these compounds (ICBG1719, ICBG1720, ICBG1724, ICBG1735, SID20373) or one or the other compound (SID20365, pyrrolnitrin; SID20345, antifungal glycopeptides). As a negative control, we also included the *Burkholderia* isolate ICBG849 as its genome does not contain either of the BGCs. The extracts were analyzed by UPLC-HRESIMS for the detection of pyrrolnitrin and antifungal glycopeptides. Pyrrolnitrin was identified by its characteristic two chlorine isotope pattern and by comparison with a pyrrolnitrin standard purchased from Millipore-Sigma. Burkholdine1213, an antifungal glycopeptide closely related to occidiofungin, was identified by comparison to the published molecular weight and molecular formula of C52H83N11O22 (36). Pyrrolnitrin was identified in ICBG1719, ICBG1720, ICBG1724, ICGB1735, SID20373, and SID20365. Burkholdine1213 was identified in ICBG1719, ICBG1720, ICBG1724, ICBG1735, SID20373, and SID20345.

After the compounds were identified in the extracts, we tested the extracts against one *Escovopsis* isolate (CF180408-01), *A. flavus*, *F. oxysporum*, and *Trichoderma* sp. in a disc diffusion assay to assess activity (Figure 4B, Figure S4). The extracts from an isolate containing both compounds (ICBG1719) demonstrated inhibition against *Escovopsis*, while extracts containing only one compound, either burkholdine or pyrrolnitrin (SID20365: pyrrolnitrin SID20345: burkholdine1213), and the DMSO control did not inhibit *Escovopsis*. This reflected the results of the previous antimicrobial inhibition bioassays using *Burkholderia* isolates. Additionally, when the extracts from SID20365 (only pyrrolnitrin) and SID20345 (only burkholdine1213) were combined, creating an extract that artificially contained both compounds, inhibition was observed (Figure 4C). This suggests that pyrrolnitrin and burkholdine 1213 may be acting additively or synergistically to inhibit *Escovopsis*.

**Figure 4.**
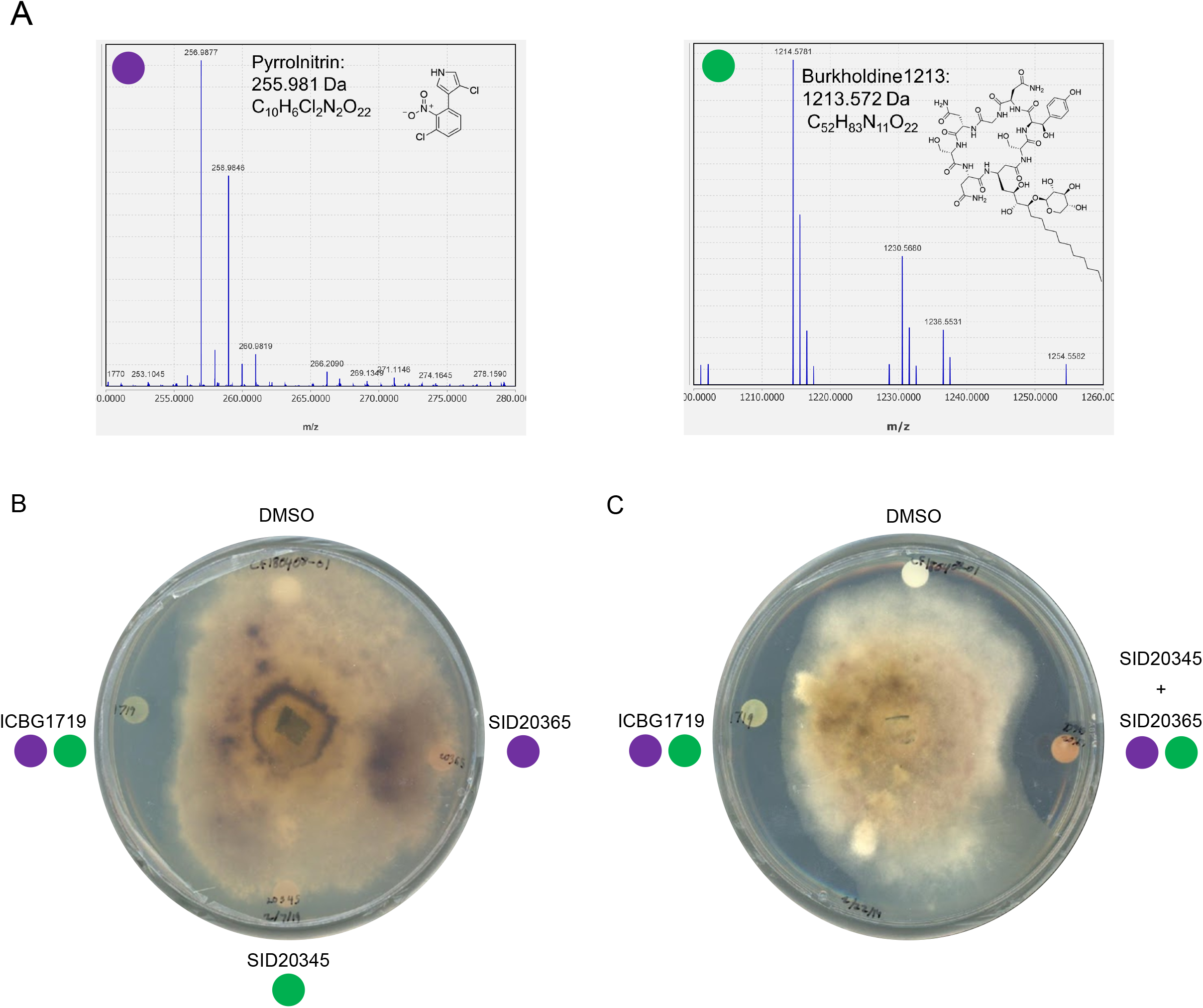
Detection of pyrrolnitrin and burkholdine1213 in organic extracts of inhibitory isolates and demonstration of extract activity. (A) Extracted Ion Chromatograms of *m/z* values that match those of pyrrolnitrin and burkholdine1213 from the organic extract of inhibitory *Burkholderia* isolate ICBG1719. (B) Disc diffusion assay of extracts from ICBG1719 (both pyrrolnitrin and burkholdine1213), SID20345 (burkholdine1213) and SID20365 (pyrrolnitrin) against *Escovopsis* sp. CF180408-01. (C) Disc diffusion assay of extracts from ICBG1719 (both pyrrolnitrin and burkholdine1213) and a combined extract of SID20345 and SID20365 (artificially containing both pyrrolnitrin and burkholdine1213) against *Escovopsis* sp. CF180408-01, demonstrating that both compounds must be present for inhibition. Additional disc diffusion assays were conducted on *Trichoderma*, *Aspergillus*, and *Fusarium* (Figure S4).

### *Burkholderia* extracts inhibit *Escovopsis* growth at lower concentrations than *Leucoagaricus* sp

The lack of inhibition of other ecologically relevant fungi suggested that the antifungals produced by inhibitory *Burkholderia* isolates may be able to inhibit *Escovopsis* while not harming *Leucoagaricus*. To test this, six *Leucoagaricus* spp. strains and an *Escovopsis* strain were grown individually on plates containing 0.005 mg/mL, 0.05 mg/mL, 0.5 mg/mL, 1 mg/mL, and 2.5 mg/mL *Burkholderia* extract from an isolate with both pyrrolnitrin and burkholdine1213 (ICBG1719). All plates with extract concentrations above 0.5 mg/mL completely inhibited all growth of *Leucoagaricus* spp. and *Escovopsis*. After six days, at 0.05mg/mL, slower growth rates (i.e., smaller diameters) were observed for four out of the six *Leucoagaricus* spp. when compared to the control, while two *Leucoagaricus* spp. and the *Escovopsis* strain demonstrated no growth (Figure S5). Finally, at 0.005 mg/mL all *Leucoagaricus* strains grew comparably to the diameter of the control, while growth of *Escovopsis* was inhibited (Figure 5).

**Figure 5.**
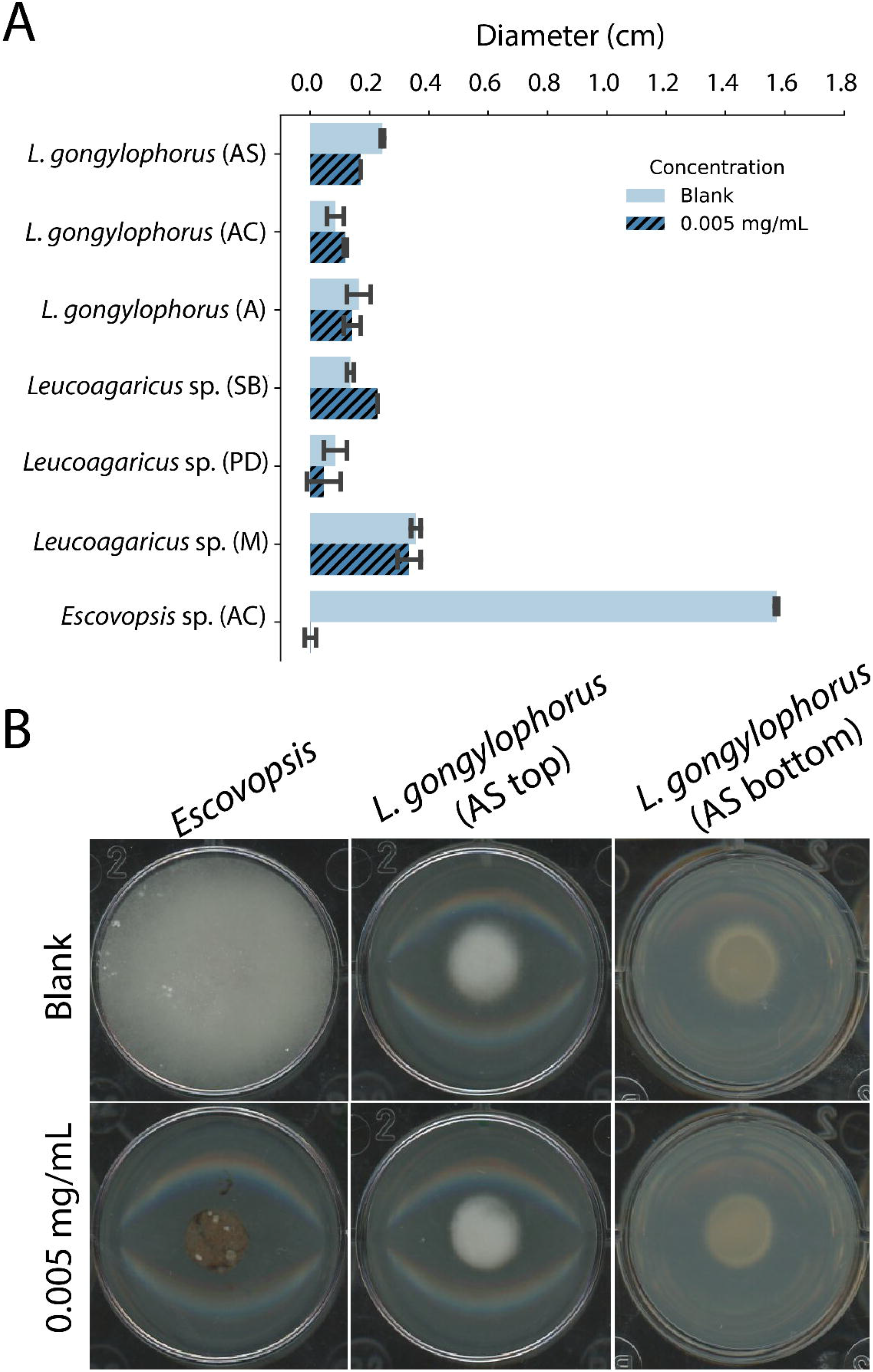
*Leucoagaricus* spp. grown on agar containing 0.005 mg/mL ICBG1719 extract (both BGCs) grows comparably to agar containing no extract, while *Escovopsis* sp. CF180408-01 from an *Atta cephalotes* colony is inhibited. Bar graph indicates the growth of *Leucoagaricus* spp. (n=2 for each strain) and *Escovopsis* (n=3) after 6 days of growth on agar containing no extract (blank) or 0.005 mg/mL (A). For visual representation, the diameter of the fungal plug (0.6mm) was subtracted from the overall diameter measurement. Photographs represent typical fungal growth of *Escovopsis* and *Leucoagaricus* spp. (photo is from *L. gongylophorus* from an *Atta sexdens* colony) on agar with no extract (blank) and 0.005 mg/mL extract (B). AS = *Atta sexdens*, AC= *Atta cephalotes*, A = *Acromyrmex* sp., SB=*Sericomyrmex bondari*, PD=*Paratrachymyrmex diversus*, M=*Myrmicocrypta* sp.

## DISCUSSION

We have demonstrated that *Burkholderia* fungus garden isolates inhibit the parasitic fungus *Escovopsis in vitro*. The inhibition of *Escovopsis* corresponds with the presence of two BGCs that encode the production of two known antifungal compounds, pyrrolnitrin and burkholdine1213. We identified both antifungals in the extracts of inhibitory isolates, confirming the expression of these clusters when cultured. A combination of the extracts of isolates that contained only one or the other antifungal did replicate the inhibitory activity, suggesting that both compounds must be present for anti-*Escovopsis* activity. Additionally, extracts of the inhibitory *Burkholderia* isolates were capable of inhibiting *Escovopsis*, but not *Leucoagaricus* spp., at concentrations as low as 0.05 mg/mL *in vitro*. These findings suggest an important role for *Burkholderia* in the defense of fungus gardens from the parasitic fungus *Escovopsis*, but not other ecologically-relevant transient fungal invaders (e.g. *Trichoderma*).

Two lineages of fungus-growing ants, *Atta* and *Sericomyrmex*, have secondarily lost the ability to harbor the defensive symbiont *Pseudonocardia* (23). Four of the five strongly inhibitory isolates were isolated from *Atta* colonies, and one isolate whose genome contained the BGC for pyrrolnitrin, SID20355, was isolated from a *Sericomyrmex* colony and had an average inhibition of 1.7 cm against the eight *Escovopsis* strains (Figure 2, Figure S2). We sampled five colonies of *Sericomyrmex* and were able to obtain at least one *Burkholderia* isolate from all five colonies, with a range of inhibition from 0.29 cm to 1.7 cm. Additional work must be done to explore whether *Burkholderia* are a potential replacement (37, 38) for *Pseudonocardia* in colonies without the symbiont. Though we equally sampled between *Pseudonocardia*-present vs *Pseudonocardia*-absent colonies (26 fungus gardens from each group), there was uneven geographic sampling and specific ant genera sampling that may influence our results (Table 1).

The presence of inhibitory *Burkholderia*, in combination with other known measures of sanitation within the fungus-growing ant system (25, 26), suggest that fungus-growing ants use multiple redundant strategies to promote healthy fungus gardens. Of note, while we focused on *Burkholderia* isolates with strong inhibition containing two BGCs, some *Burkholderia* isolates from fungus gardens containing either one BGC or neither were still able to less severely inhibit *Escovopsis* (Figure 2, Figure S2). This indicates that harboring *Burkholderia* that can inhibit *Escovopsis* to some degree may be a strategy used across the ant phylogeny. However, this strategy cannot be used with other ecologically relevant fungi, as we saw *Burkholderia* isolates barely, if at all, inhibited *Trichoderma, Aspergillus, and Fusarium*. However, other work with different fungus garden isolates have indicated inhibition of more transient fungi, including *Trichoderma* (27, 39).

Similarly to internal animal gut microbiomes, the fungus garden microbiome is composed of a complex and diverse microbial community where the roles of resident bacteria are still being discovered. In recent years, there has been an accumulation of evidence that members of the fungus garden microbiome potentially function as beneficial symbionts, with some taxa aiding in nitrogen fixation, degradation of plant matter, and/or detoxification of plant secondary compounds. In this study, we have described a fungus garden associated bacterial genus, *Burkholderia* spp., as a possible mechanism for certain fungus-growing ant lineages to defend their fungus gardens from parasites. Overall, these new findings combined with past studies suggests that members of the fungus garden microbiome play a key role in facilitating the response of fungus-growing ants to environmental changes and pressures.

## MATERIALS AND METHODS

### Sampling and bacterial isolations

We collected fungus garden samples in January 2017 and April 2018 in the following locations: Anavilhanas, Amazonas, Brazil; Ducke Reserve, Amazonas, Brazil; Itatiaia, Rio de Janeiro, Brazil; Sao Paulo State, Brazil and La Selva Biological Station, Costa Rica (Table S1). Collections of biological samples and research on genetic resources were authorized in Brazil by SISBIO #46555–5 and CNPq #010936/2014–9. In Costa Rica, permits were granted by Comisión Institucional de Biodiversidad of the University of Costa Rica (Resolucion no. 009), the Organization for Tropical Studies (OTS), and the Ministerio de Ambiente y Energía (MINAE). For the fungus garden samples collected in Brazil, for each colony, a small fungus garden piece collected from the inner region of the garden was put into PBS and then plated onto Yeast Malt Extract Agar (YMEA). Plates with bacterial colonies were brought back to the UW-Madison and pure bacterial isolates were obtained after several rounds of subculturing. We obtained a total of 317 bacterial isolates. We identified 117 isolates to genus-level by 16S rRNA sequencing. Of those, 33 corresponded to the genus *Burkholderia*. For the fungus garden samples collected in Costa Rica, ~0.2 g of fungus garden were collected from the inner region of the garden, put into PBS, vortexed and homogenized, then the PBS-fungus garden homogenate was serially diluted and all dilutions were plated on both YMEA and modified APCA (40) (per L: 0.79 g (NH_4_)_2_SO_4_, 1 g KH_2_PO_4_, 0.5 g MgSO_4_ - &H2O, 0.2 g KCl, 2 g L-Arabinose, 5 mg Crystal Violet, 50 mg polymixin B sulphate, 50 mg ampicillin sodium, 10 mg chloramphenicol, 15 g agar).

### DNA Extraction and Assembly

We extracted DNA from 30 bacterial isolates using the Wizard Genomic DNA Purification Kit (Promega, USA) using the Gram-Negative Protocol. Genomic DNA libraries for Illumina MiSeq 2×300bp paired-end sequencing were prepared by the University of Wisconsin - Madison Biotechnology Center. Reads were corrected with MUSKETv1.1 (41), paired-ends were merged with FLASH v.1.2.7 (42), and assembled with SPAdes 3.11.0 (43). We taxonomically classified the *Burkholderiaceae* isolates to species-level by performing a Tetra Correlation Search at JSpecies WS (44). In addition, ANI was calculated using the anvi’o (45) anvi-compute-genome-similarity program, with the default pyani (46) settings.

### Phylogenetic Tree

We generated a genome-based, multilocus *Burkholderiaceae* phylogeny based on previous methods (47). Briefly, the phylogeny was generated using 93 full TIGRFAM proteins in the “core bacterial protein” set (GenProp0799) as the molecular data set. The protein sequences with the top HMMer bitscore for each protein family were aligned using MAFFT (48) which were then converted to codon alignments and concatenated. RAxML-7.2.6 (49)was used to generate phylogeny using the GTRgamma substitution model and 100 rapid bootstraps on the final, recombination-free alignment. The gene tree-based phylogeny was generated using ASTRAL-II (50). Phylogenies were visualized and edited in FigTree v. 1.4.3.A. We constructed the 16S rRNA phylogenetic tree using BEAST2 v2.5.1 (51) with a TN93 substitution model. The analysis was run for 150,000,000 generations, sampling every 1,000 generations, and a burn-in of 10% was applied.

### *Escovopsis*-*Burkholderiaceae* bioassays

In order to assess inhibition profiles of *Burkholderiaceae* against *Escovopsis* and other fungi, we performed *in vitro* plate assays. For each isolate, we grew an overnight culture in Yeast Malt Extract (YME) for 16-24 hours. We spotted 10 μL of *Burkholderiaceae* onto the middle of a 100 mm Potato-Dextrose Agar (PDA) plate and incubated at room temperature for seven days. After seven days, 6mm fungal plugs of *Escovopsis*, *Trichoderma, Aspergillus,* and *Fusarium* were put on the edge of the plate. Control plates containing only a *Burkholderiaceae* isolate or only a fungus were also made for each isolate. Over the course of a month both sides of the plate were scanned every 6-7 days and the zone of inhibition was measured using Fiji (52).

### Biosynthetic Gene Cluster Annotation

We used antiSMASH v4.0 (53) to predict BGCs and BiG-SCAPE (54) to construct sequence similarly networks of BGCs. In addition to our predicted BGCs for the BiG-SCAPE analysis, we included the reference BGCs from the MiBiG repository. We used cytoscape (55) to visualize the BGC networks. We color-coded the clusters into two categories: inhibitory *Burkholderia* isolates (average ZOI ≥ 2.45cm against *Escovopsis*) and non-inhibitory *Burkholderia* isolates (average ZOI ≤ 2.45cm against *Escovopsis*) and looked for clusters that contained all the inhibitory isolates.

### *Burkholderia* extracts

*Burkholderia* isolates ICBG1719, ICBG1720, ICBG1724, ICGB1735, SID20373, SID20345, SID20365, and ICBG849 were inoculated from YMEA plates into 100 mL of YME broth in 500 mL baffled flasks and shaken for 36 hours. The 100 mL cultures were used to inoculate two 500 mL cultures of YME broth in 2 L flasks for each isolate. The isolates were shaken for 36 hours with Diaion HP20 resin. The cultures were then vacuum filtered through a Whatman 1 sized filter paper and the Diaion resin and cell mass was extracted overnight with ethyl acetate. Excess sodium sulfate anhydrous was added to the extraction to remove residual water. The ethyl acetate was filtered and dried in vacuo to yield a *Burkholderia* extract.

### HRESIMS of *Burkholderia* extracts

*Burkholderia* extracts were resuspended in methanol and analyzed for the presence of pyrrolnitrin and burkholdine1213 by UPLC-HRESIMS on a Q Exactive Orbitrap mass spectrometer. A liquid chromatography gradient was run from 5% acetonitrile with 0.1% formic acid to 100% acetonitrile with 0.1% formic acid over 15 minutes on a Phenomenex XB C18, 2.1 mm by 100 mm, 2.6 μm particle size column. The scan range was from 200 *m/z* to 2,000 *m/z* in positive mode.

### *Escovopsis* - *Burkholderia* extract assays

*Escovopsis* strain ICBG1053 (*Apterostigma*) and CF180408-01 (*Atta cephalotes*) plugs were plated onto PDA and grown for three days until white mycelia could be seen. Additionally, *Trichoderma*, *Aspergillus*, and *Fusarium* were plated on PDA and grown until slight mycelial growth was visible. *Burkholderia* extracts from ICBG1719, SID20345, and SID20365 were dissolved in dimethyl sulfoxide (DMSO). The extracts were pipetted onto sterile filter paper discs in 10μL of DMSO at 0.5 mg/disc, 1 mg/disc, and 2 mg/disc and placed onto the PDA plates containing the *Escovopsis* or other fungi along with a DMSO control disc. After two weeks of growth the ability of each extract to inhibit the growth of the fungi was assessed and pictures were taken of the plates.

### *Leucoagaricus-Burkholderia* extract assays

We plated six *Leucoagaricus* strains isolated from *Atta sexdens, Atta cephalotes, Acromyrmex octospinosus, Sericomyrmex bondari, Paratrachymyrmex diversus, Myrmicocrypta* sp. and an *Escovopsis* isolate from *Atta cephalotes* on PDA and let them grow for a month at room temperature. We prepared PDA plates containing 0 mg/mL, 0.005 mg/mL, 0.05 mg/mL, 0.5 mg/mL, 1 mg/mL, and 2.5 mg/mL *Burkholderia* extract in DMSO. Media was vigorously mixed for homogenous distribution of the extract, and then 3 mL was pipetted into each well of a 12-well plate (manufacturing details) and left to dry overnight in the dark. Then, 0.6 mm plugs taken from the outer edges of *Leucoagaricus* spp. or *Escovopsis* fungal plates were deposited into the center of each well (n=2 for each fungal strain × 6 concentrations). Pictures were taken 6 days post-exposure and the diameter of fungal growth was measured in Fiji.

## Data availability

All sequencing data have been uploaded to NCBI under the following BioProject numbers: PRJNA564151 and PRJNA603049. Whole genome and SRA accession numbers for each isolate can be found in Table S2.

## Acknowledgements

We are grateful to Ted Schultz, Jeffrey Sosa-Calvo, and Eugenia Okonski for identification of our fungus-growing ant specimens. We thank Dr. Camila Carlos-Shanley for her guidance and support at the conception of this project. We thank the staff and scientists at the La Selva Biological Station for allowing field collections and the scientists Allan Artavia and Miguel Pacheco in the Pinto-Tomas lab at the University of Costa Rica for their assistance with field collections.

## Funding

This material is based upon work supported in part by the National Institutes of Health grant U19 TW009872, National Institutes of Health grant U19 AI142720, the National Institute of Allergy and Infectious Diseases of the National Institutes of Health grant T32 AI055397, National Science Foundation grant DEB-1927155, and the São Paulo Research Foundation (FAPESP) grant ##2013/50954-0. The content is solely the responsibility of the authors and does not necessarily represent the official views of the National Institutes of Health.

---

**Table S1.** Collection information for fungal and bacterial isolates

**Table S2.** Genome statistics and accession numbers for 30 bacterial isolates

**Table S3.** Post-hoc pairwise Student’s *t*-test results for the zones of inhibition of *Escovopsis* spp. by all 32 *Burkholderiaceae* isolates.

**Dataset S1.** 16S rRNA sequences without accession numbers used in this study

**Dataset S2.** Average Nucleotide Identity matrix

**Figure S1**. 16S rRNA (A) and whole genome (B) phylogenies of *Burkholderiaceae* isolates.

**Figure S2**. Zones of inhibition of *Escovopsis* spp. by all 32 *Burkholderiaceae* isolates. Red boxplots indicate isolates with significantly higher ZOIs (ZOI ≥ 2.45 cm). Blue boxplots have significantly lower ZOIs (ZOIs < 2.45 cm). A one-way ANOVA test [F(31,192)=24.41, p<0.0001] was used to assess significance between strongly inhibitory and less inhibitory isolates. The post-hoc Student’s *t*-test between all pairings are provided in Dataset S2.

**Figure S3.** 13 additional *Burkholderiaceae* isolates from *Sericomyrmex amabilis* colonies were tested against *Escovopsis* spp. Hatched squares indicate the pairings that could not be measured due to various contamination and growth complications.

**Figure S4.** Disc diffusion assay of extracts from ICBG1719 (both pyrrolnitrin and burkholdine1213) and a combined extract of SID20345 and SID20365 (artificially containing both pyrrolnitrin and burkholdine1213) against (A) *Aspergillus flavus* (B) *Fusarium oxysporum* and (C) *Trichoderma* sp.

**Figure S5.** *Leucoagaricus* spp. and *Escovopsis* sp. CF180408-01 grown on agar containing 0.05 mg/mL ICBG1719 extract (both BGCs). Bar graph indicates the growth of *Leucoagaricus* spp. (n=2 for each strain) and *Escovopsis* (n=3) after 6 days of growth on agar containing no extract (blank) or 0.05 mg/mL. AS = *Atta sexdens*, AC= *Atta cephalotes*, A = *Acromyrmex* sp., SB=*Sericomyrmex bondari*, PD=*Paratrachymyrmex diversus*, M=*Myrmicocrypta* sp.

## Notes

### Competing Interest Statement

The authors have declared no competing interest.

## REFERENCES

1. Skelton J, Doak S, Leonard M, Creed RP, Brown BL. 2016. The rules for symbiont community assembly change along a mutualism – parasitism continuum 843–853.

2. Douglas AE. 1998. Nutritional interactions in insect-microbial symbioses: Aphids and their symbiotic bacteria Buchnera. Annu Rev Entomol 43:17–37.

3. Douglas AE. 2009. The microbial dimension in insect nutritional ecology. Funct Ecol 23:38–47.

4. Flórez L V., Biedermann PHW, Engl T, Kaltenpoth M. 2015. Defensive symbioses of animals with prokaryotic and eukaryotic microorganisms. Nat Prod Rep 32:904–936.

5. Van Arnam EB, Currie CR, Clardy J. 2018. Defense contracts: Molecular protection in insect-microbe symbioses. Chem Soc Rev 47:1638–1651.

6. Dillon RJ, Dillon VM. 2004. The gut bacteria of insects: Nonpathogenic Interactions. Annu Rev Entomol 49:71–92.

7. Piel J. 2009. Metabolites from symbiotic bacteria. Nat Prod Rep 26:338–362.

8. Pan X, Zhou G, Wu J, Bian G, Lu P, Raikhel AS, Xi Z. 2012. Wolbachia induces reactive oxygen species (ROS)-dependent activation of the Toll pathway to control dengue virus in the mosquito Aedes aegypti. Proc Natl Acad Sci U S A 109.

9. Solomon SE, Rabeling C, Sosa-Calvo J, Lopes CT, Rodrigues A, Vasconcelos HL, Bacci M, Mueller UG, Schultz TR. 2019. The molecular phylogenetics of Trachymyrmex Forel ants and their fungal cultivars provide insights into the origin and coevolutionary history of ‘higher-attine’ ant agriculture. Syst Entomol 44:939–956.

10. Sosa-Calvo J, Branstetter MG, Jes A, Brady G, Schultz TR, Lloyd MW, Faircloth BC, Mw L, Bc F, Sg B, Branstetter MG, Schultz TR, Ješovnik A, Sosa-calvo J, Lloyd MW, Faircloth BC, Brady SG, Schultz TR. 2017. Dry habitats were crucibles of domestication in the evolution of agriculture in ants. Proc R Soc B Biol Sci 284.

11. Currie CR. 2001. A Community of Ants, Fungi, and Bacteria: A Multilateral Approach to Studying Symbiosis. Annu Rev Microbiol 55:357–380.

12. Aylward FO, Currie CR, Suen G. 2012. The evolutionary innovation of nutritional symbioses in leaf-cutter ants. Insects 3:41–61.

13. Khadempour L, Burnum-Johnson KE, Baker ES, Nicora CD, Webb-Robertson B-JM, White RA, Monroe ME, Huang EL, Smith RD, Currie CR. 2016. The fungal cultivar of leaf-cutter ants produces specific enzymes in response to 2 different plant substrates 1–51.

14. Aylward FO, Burnum KE, Scott JJ, Suen G, Tringe SG, Adams SM, Barry KW, Nicora CD, Piehowski PD, Purvine SO, Starrett GJ, Goodwin LA, Smith RD, Lipton MS, Currie CR. 2012. Metagenomic and metaproteomic insights into bacterial communities in leaf-cutter ant fungus gardens. ISME J 6:1688–1701.

15. Suen G, Scott JJ, Aylward FO, Adams SM, Tringe SG, Pinto-Tomás AA, Foster CE, Pauly M, Weimer PJ, Barry KW, Goodwin LA, Bouffard P, Li L, Osterberger J, Harkins TT, Slater SC, Donohue TJ, Currie CR. 2010. An insect herbivore microbiome with high plant biomass-degrading capacity. PLoS Genet 6.

16. Khadempour L, Fan H, Keefover-Ring K, Carlos-Shanley C, Nagamoto NS, Dam MA, Pupo MT, Currie CR. 2020. Metagenomics Reveals Diet-Specific Specialization of Bacterial Communities in Fungus Gardens of Grass- and Dicot-Cutter Ants. Front Microbiol 11:1–14.

17. Barcoto MO, Carlos-Shanley C, Fan H, Ferro M, Nagamoto NS, Bacci M, Currie CR, Rodrigues A. 2020. Fungus-growing insects host a distinctive microbiota apparently adapted to the fungiculture environment. Sci Rep 10:1–13.

18. Ronque MUV, Lyra ML, Migliorini GH, Bacci M, Oliveira PS. 2020. Symbiotic bacterial communities in rainforest fungus-farming ants: evidence for species and colony specificity. Sci Rep 10:1–12.

19. Francoeur C, Khadempour L, Moreira-Soto R, Gotting K, Book A, Pinto-Tomás A, Keefover-Ring K, Currie C. 2020. Bacteria contribute to plant secondary compound degradation in a generalist herbivore system. MBio 11:1–18.

20. Pinto-Tomás AA, Anderson MA, Suen G, Stevenson DM, Chu FST, Cleland WW, Weimer PJ, Currie CR. 2009. Symbiotic nitrogen fixation in the fungus gardens of leaf-cutter ants. Science (80−) 326:1120–1123.

21. Currie CR, Mueller UG, Malloch D. 1999. The agricultural pathology of ant fungus gardens. Proc Natl Acad Sci U S A 96:7998–8002.

22. Currie CR, Poulsen M, Mendenhall J, Boomsma JJ, Billen J. 2006. Coevolved crypts and exocrine glands support mutualistic bacteria in fungus-growing ants. Science (80−) 311:81–83.

23. Li H, Sosa-Calvo J, Horn HA, Pupo MT, Clardy J, Rabeling C, Schultz TR. 2018.Convergent evolution of complex structures for ant-bacterial defensive symbiosis in fungus-farming ants [Evolution]. Proc Natl Acad Sci U S A 1–6.

24. Currie CR, Summerbell RC, Scott JA, Malloch D. 1999. Fungus-growing ants use antibiotic-producing bacteria to control garden parasites. Nature 398:701–704.

25. Currie CR, Stuart AE. 2001. Weeding and grooming of pathogens in agriculture by ants. Proc R Soc B Biol Sci 268:1033–1039.

26. Fernández-Marín H, Nash DR, Higginbotham S, Estrada C, Van Zweden JS, D’Ettorre P, Wcislo WT, Boomsma JJ. 2015. Functional role of phenylacetic acid from metapleural gland secretions in controlling fungal pathogens in evolutionarily derived leaf-cutting ants. Proc R Soc London B 282:2–12.

27. Santos AV, Dillon RJ, Dillon VM, Reynolds SE, Samuels RI. 2004. Ocurrence of the antibiotic producing bacterium Burkholderia sp. in colonies of the leaf-cutting ant Atta sexdens rubropilosa. FEMS Microbiol Lett 239:319–323.

28. Kaltenpoth M, Flórez L V. 2020. Versatile and dynamic symbioses between insects and burkholderia bacteria. Annu Rev Entomol 65:145–170.

29. Partida-Martinez LP, Hertweck C. 2005. Pathogenic fungus harbours endosymbiotic bacteria for toxin production. Nature 437:884–888.

30. Partida-Martinez LP, Groth I, Schmitt I, Richter W, Roth M, Hertweck C. 2007. Burkholderia rhizoxinica sp. nov. and Burkholderia endofungorum sp. nov., bacterial endosymbionts of the plant-pathogenic fungus Rhizopus microsporous. Int J Syst Evol Microbiol 57:2583–2590.

31. Flórez L V., Scherlach K, Gaube P, Ross C, Sitte E, Hermes C, Rodrigues A, Hertweck C, Kaltenpoth M. 2017. Antibiotic-producing symbionts dynamically transition between plant pathogenicity and insect-defensive mutualism. Nat Commun 8.

32. Flórez L V., Scherlach K, Miller IJ, Rodrigues A, Kwan JC, Hertweck C, Kaltenpoth M. 2018. An antifungal polyketide associated with horizontally acquired genes supports symbiont-mediated defense in Lagria villosa beetles. Nat Commun 9.

33. Jung BK, Hong SJ, Park GS, Kim MC, Shin JH. 2018. Isolation of Burkholderia cepacia JBK9 with plant growth-promoting activity while producing pyrrolnitrin antagonistic to plant fungal diseases. Appl Biol Chem 61:173–180.

34. Hammer PE, Hill DS, Lam ST, Van Pée KH, Ligon JM. 1997. Four genes from Pseudomonas fluorescens that encode the biosynthesis of pyrrolnitrin. Appl Environ Microbiol 63:2147–2154.

35. Gu G, Smith L, Liu A, Lu SE. 2011. Genetic and biochemical map for the biosynthesis of occidiofungin, an antifungal produced by Burkholderia contaminans strain MS14. Appl Environ Microbiol 77:6189–6198.

36. Lin Z, Falkinham JO, Tawfik KA, Jeffs P, Bray B, Dubay G, Cox JE, Schmidt EW. 2012. Burkholdines from Burkholderia ambifaria: Antifungal agents and possible virulence factors. J Nat Prod 75:1518–1523.

37. Husnik F, McCutcheon JP. 2016. Repeated replacement of an intrabacterial symbiont in the tripartite nested mealybug symbiosis. Proc Natl Acad Sci U S A 113:E5416–E5424.

38. Sudakaran S, Kost C, Kaltenpoth M. 2017. Symbiont Acquisition and Replacement as a Source of Ecological Innovation. Trends Microbiol 25:375–390.

39. Schoenian I, Spiteller M, Ghaste M, Wirth R, Herz H, Spiteller D. 2011. Chemical basis of the synergism and antagonism in microbial communities in the nests of leaf-cutting ants. Proc Natl Acad Sci U S A 108:1955–1960.

40. Kawanishi T, Uematsu S, Nishimura K, Otani T, Tanaka-Miwa C, Hamamoto H, Namba S. 2009. A new selective medium for Burkholderia caryophylli, the causal agent of carnation bacterial wilt. Plant Pathol 58:237–242.

41. Liu Y, Schröder J, Schmidt B. 2013. Musket: A multistage k-mer spectrum-based error corrector for Illumina sequence data. Bioinformatics 29:308–315.

42. Magoč T, Salzberg SL. 2011. FLASH: Fast length adjustment of short reads to improve genome assemblies. Bioinformatics 27:2957–2963.

43. Bankevich A, Nurk S, Antipov D, Gurevich AA, Dvorkin M, Kulikov AS, Lesin VM, Nikolenko SI, Pham S, Prjibelski AD, Pyshkin A V., Sirotkin A V., Vyahhi N, Tesler G, Alekseyev MA, Pevzner PA. 2012. SPAdes: A new genome assembly algorithm and its applications to single-cell sequencing. J Comput Biol 19:455–477.

44. Richter M, Rosselló-Móra R, Oliver Glöckner F, Peplies J. 2016. JSpeciesWS: A web server for prokaryotic species circumscription based on pairwise genome comparison. Bioinformatics 32:929–931.

45. Eren AM, Esen OC, Quince C, Vineis JH, Morrison HG, Sogin ML, Delmont TO. 2015. Anvi’o: An advanced analysis and visualization platformfor’omics data. PeerJ 2015:1–29.

46. Pritchard L, Glover RH, Humphris S, Elphinstone JG, Toth IK. 2016. Genomics and taxonomy in diagnostics for food security: Soft-rotting enterobacterial plant pathogens. Anal Methods 8:12–24.

47. McDonald BR, Currie CR. 2017. Lateral Gene Transfer Dynamics in the Ancient Bacterial Genus Streptomyces. MBio 8:1–12.

48. Katoh K, Standley DM. 2013. MAFFT multiple sequence alignment software version 7: Improvements in performance and usability. Mol Biol Evol 30:772–780.

49. Stamatakis A. 2006. RAxML-VI-HPC: maximum likelihood-based phylogenetic analyses with thousands of taxa and mixed models. Bioinformatics 22:2688–2690.

50. Mirarab S, Warnow T. 2015. ASTRAL-II: Coalescent-based species tree estimation with many hundreds of taxa and thousands of genes. Bioinformatics 31:i44–i52.

51. Bouckaert R, Vaughan TG, Barido-Sottani J, Duchêne S, Fourment M, Gavryushkina A, Heled J, Jones G, Kühnert D, De Maio N, Matschiner M, Mendes FK, Müller NF, Ogilvie HA, Du Plessis L, Popinga A, Rambaut A, Rasmussen D, Siveroni I, Suchard MA, Wu CH, Xie D, Zhang C, Stadler T, Drummond AJ. 2019. BEAST 2.5: An advanced software platform for Bayesian evolutionary analysis. PLoS Comput Biol 15:1–28.

52. Schindelin J, Arganda-Carreras I, Frise E, Kaynig V, Longair M, Pietzsch T, Preibisch S, Rueden C, Saalfeld S, Schmid B, Tinevez JY, White DJ, Hartenstein V, Eliceiri K, Tomancak P, Cardona A. 2012. Fiji: An open-source platform for biological-image analysis. Nat Methods 9:676–682.

53. Blin K, Wolf T, Chevrette MG, Lu X, Schwalen CJ, Kautsar SA, Suarez Duran HG, De Los Santos ELC, Kim HU, Nave M, Dickschat JS, Mitchell DA, Shelest E, Breitling R, Takano E, Lee SY, Weber T, Medema MH. 2017. AntiSMASH 4.0 - improvements in chemistry prediction and gene cluster boundary identification. Nucleic Acids Res 45:W36–W41.

54. Navarro-Muñoz JC, Selem-Mojica N, Mullowney MW, Kautsar SA, Tryon JH, Parkinson EI, De Los Santos ELC, Yeong M, Cruz-Morales P, Abubucker S, Roeters A, Lokhorst W, Fernandez-Guerra A, Cappelini LTD, Goering AW, Thomson RJ, Metcalf WW, Kelleher NL, Barona-Gomez F, Medema MH. 2020. A computational framework to explore large-scale biosynthetic diversity. Nat Chem Biol 16:60–68.

55. Shannon P, Markiel A, Ozier O, Baliga NS, Wang JT, Ramage D, Amin N, Schwikowski B, Ideker T. 2003. Cytoscape: A Software Environment for Integrated Models. Genome Res 13:2498–2504.

