## Supplementary figures and images for "*Burkholderia* from fungus gardens of fungus-growing ants produce antifungals that inhibit the specialized parasite *Escovopsis*"

### Supplemental Figure 1

A 16S rRNA phylogenetic tree

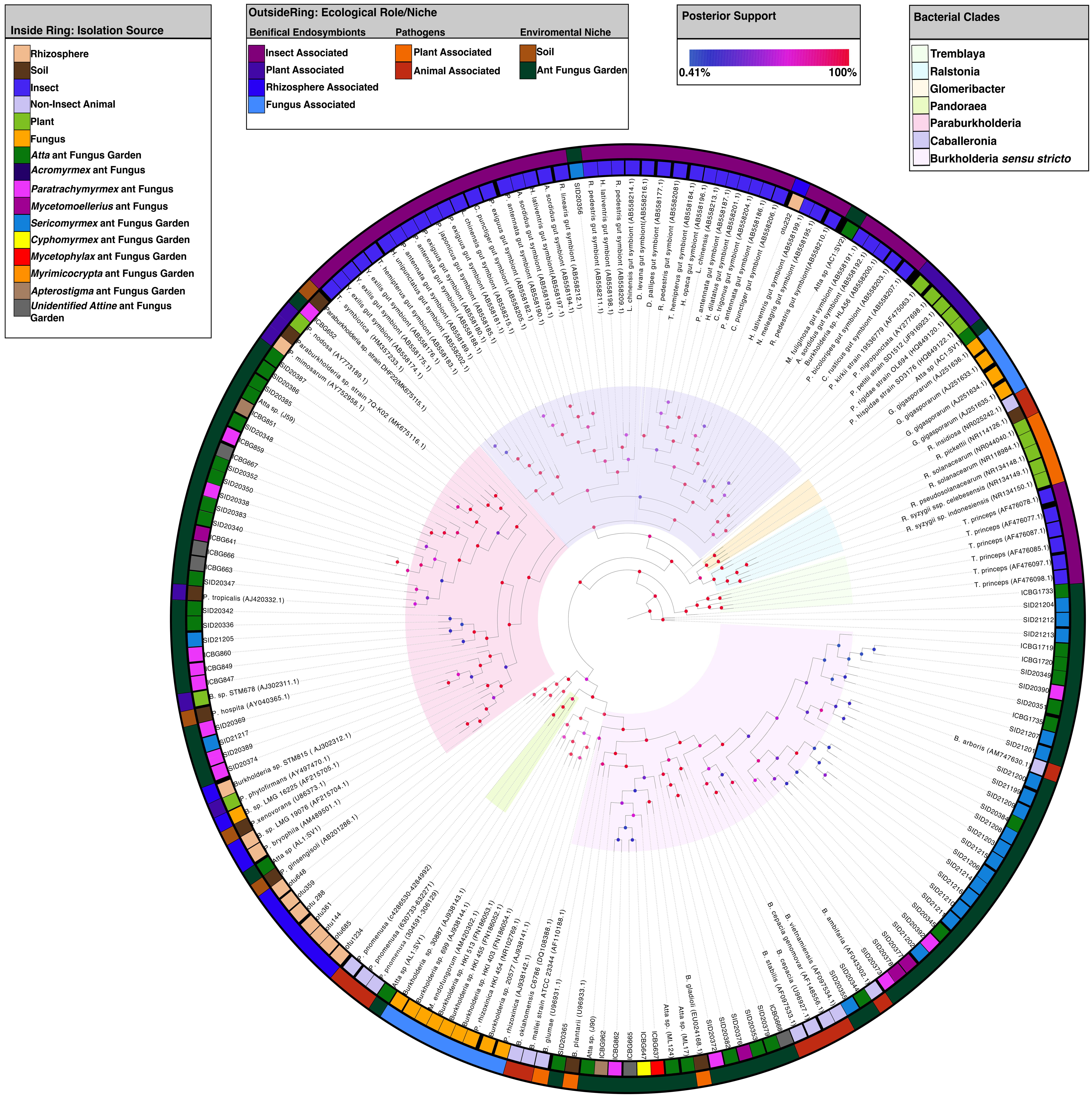


B Whole genome phylogenetic tree

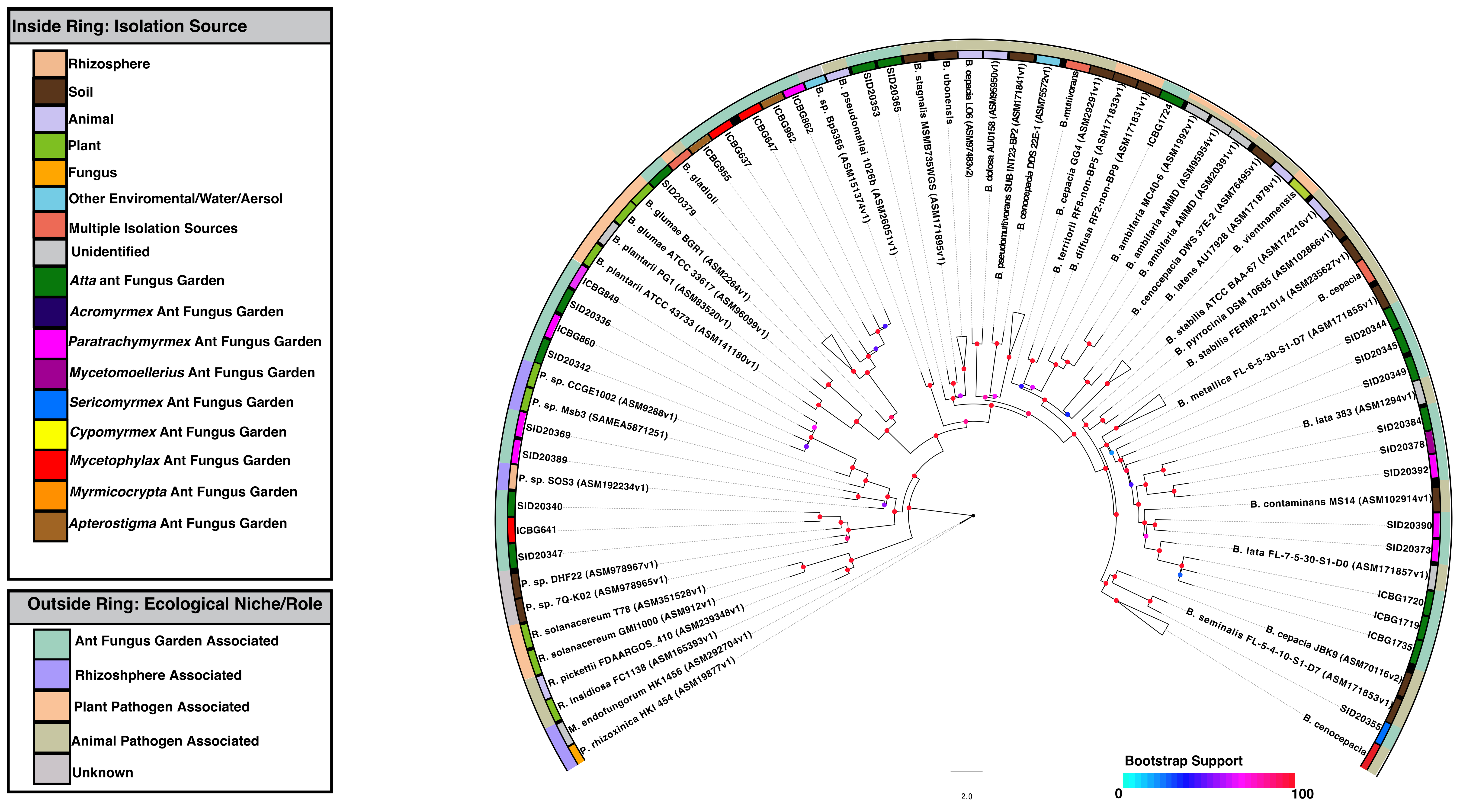

### Supplemental Figure 2

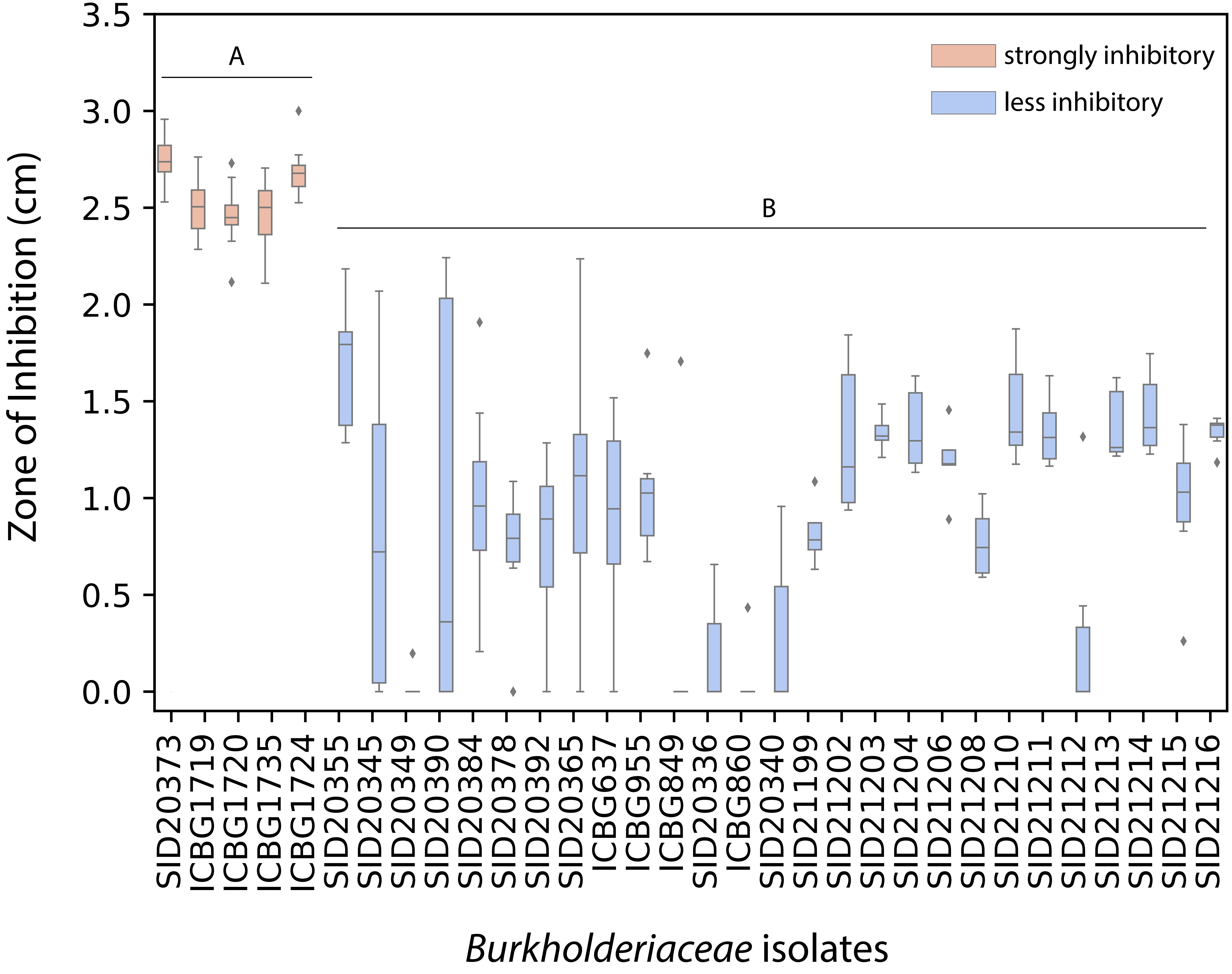

### Supplemental Figure 3

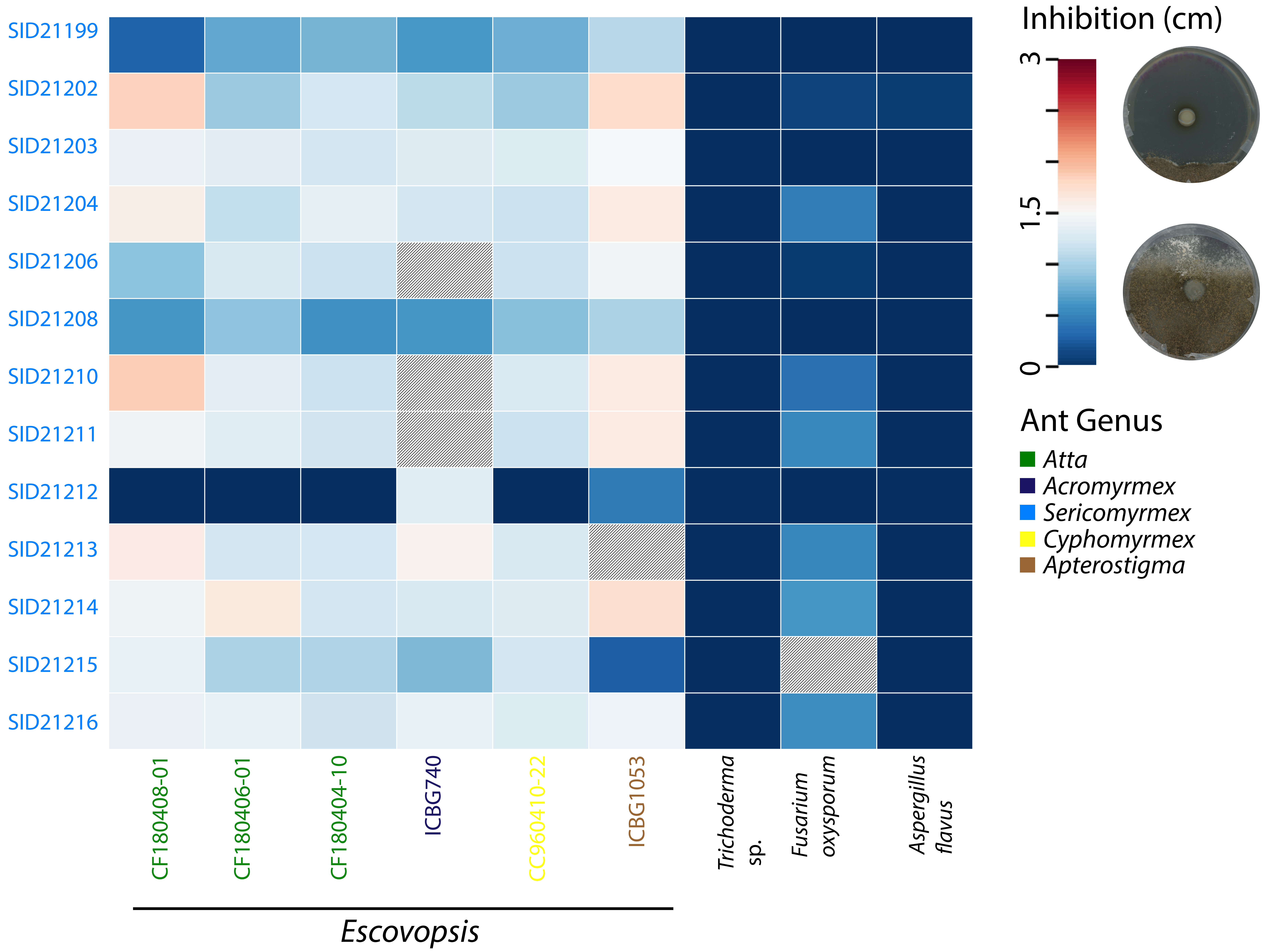

### Supplemental Figure 4

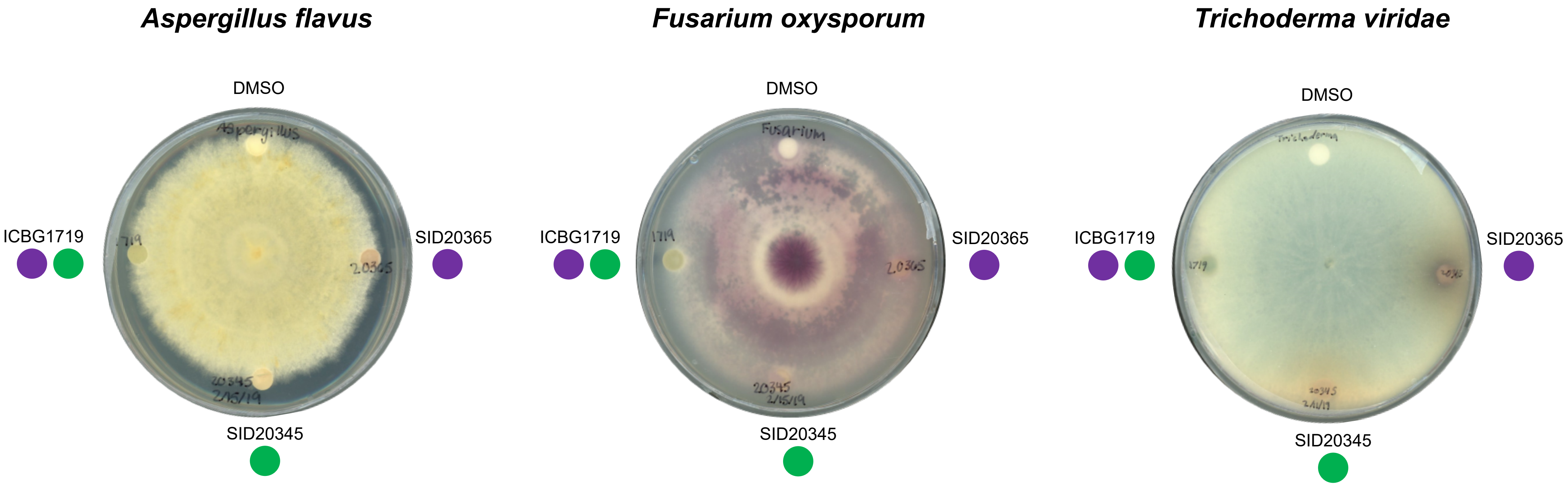

### Supplemental Figure 5

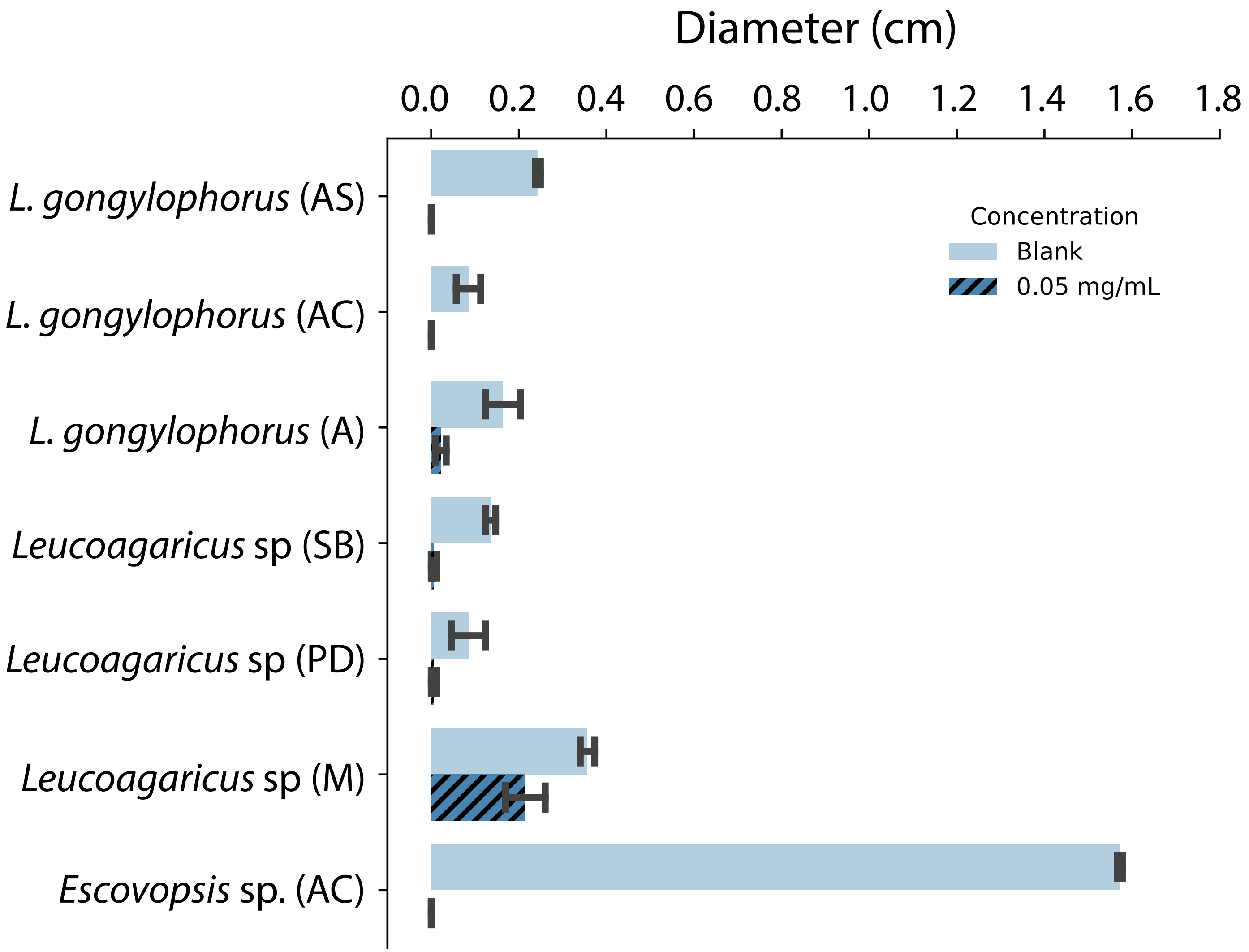
