## Supplemental Table 1 for "*Burkholderia* from fungus gardens of fungus-growing ants produce antifungals that inhibit the specialized parasite *Escovopsis*"

| Isolate ID | Genus (16S) | Host colony | Host-ID | GPS Coordinates | Location | Notes | Date collected (m/d/y) | colony code |
|---|---|---|---|---|---|---|---|---|
| ICBG2046 | Escovopsis | Cyphomyrmex longiscapus | n/a | n/a | Panama | unidentified species | 4/10/96 | CC960410-22 |
| ICBG2049 | Escovopsis | Myrmicocrypta | n/a | n/a | Panama | species ID: Escovopsis kreiselii | 4/15/96 | CC960415-06 |
| ICBG740 | Escovopsis | Acromyrmex | 2160 | n/a | Anavilhanas, AM | species ID: Escovopsis weberi | 1/21/17 | CAR170121-01 |
| ICBG1053 | Escovopsis | Apterostigma | 2139 | S2° 31.430 W60° 49.477 | Anavilhanas, AM | species ID: Escovopsis clavatus | 1/20/17 | AR170120-08 |
| CF180404-10 | Escovopsis | Atta cephalotes | n/a | N10.43311, W84.01439 | La Selva Biological Station, CR | species ID: Escovopsis weberi | 4/4/18 | CF180404-10 |
| CF180408-01 | Escovopsis | Atta cephalotes | n/a | n/a | La Selva Biological Station, CR | species ID: Escovopsis weberi | 4/8/18 | CF180408-01 |
| CF180406-01 | Escovopsis | Atta cephalotes | n/a | N10.42764, W84.00176 | La Selva Biological Station, CR | species ID: Escovopsis weberi | 4/6/18 | CF180406-01 |
| HH180403-03 | Escovopsis | Atta cephalotes | n/a | N10.43136, W084.00557 | La Selva Biological Station, CR | species ID: Escovopsis weberi | 4/3/18 | HH180403-03 |
| Trichoderma sp. | | N/A | PID44 | | | | | |
| Aspergillus flavus | | Scale insect | PID14 | | | NRRL 485, ATCC 11492 | | |
| Fusarium oxysporum | | N/A | n/a | | | subspecies: f. sp. lycopersici 4287 | | |
| AS2 | Leucoagaricus | Atta sexdens | n/a | #-21.15406, -47.8571 | Sao Paulo, Brazil | Leucoagaricus gongylophorus | 11/29/14 | LK141129-01 |
| CF190323-01 | Leucoagaricus | Atta cephalotes | n/a | N10.42746, W084.00158 | La Selva Biological Station, CR | Leucoagaricus gongylophorus | 3/23/19 | CF190323-01 |
| Acro colony 3 | Leucoagaricus | Acromyrmex | n/a | n/a | University of Costa Rica campus | Leucoagaricus gongylophorus | 3/12/19 | AA190312-01 |
| AJ141001-05 | Leucoagaricus | Sericomyrmex bondari | n/a | #-6.063, -50.057 | Pará, Parauapebas, FL Nacional de Carajás | | | |
| WM170124-07 | Leucoagaricus | Paratrachymyrmex diversus | 2255 | S2° 16' 13.7 W61° 01' 05.3 | Anavilhanas, AM | | 1/24/17 | WM170124-07 |
| LK170123-01 | Leucoagaricus | Myrmicocrypta | 2186 | S2° 35.9 W61° 1.50 | Anavilhanas, AM | | 1/23/17 | LK170123-01 |
| ICBG637 | Burkholderia | Mycetophylax cf. plaumanni | 2324 | S2° 55.808 W59° 58.479 | Ducke Reserve, AM | | 1/28/17 | TF170128-03 |
| ICBG647 | Burkholderia | Cyphomyrmex | 2320 | S2° 55.808 W59° 58.479 | Ducke Reserve, AM | | 1/28/17 | WM170128-05 |
| ICBG1735 | Burkholderia | Atta sp. | 2341 | -22.84652, -48.43412 | Sao Paulo State | | 1/6/17 | LK170106-06 |
| ICBG862 | Burkholderia | Paratrachymyrmex | 2256 | S2° 16' 13.7 W61° 01' 05.3 | Anavilhanas, AM | | 1/24/17 | WM170124-08 |
| ICBG955 | Burkholderia | Apterostigma sp. | 2086 | S2° 32' 1.9 W60° 50' 6.1 | Anavilhanas, AM | | 1/19/17 | BP170119-02 |
| ICBG962 | Burkholderia | Apterostigma sp. | 2149 | S2° 31' 23,4 W60° 49' 31,9 | Anavilhanas, AM | | 1/20/17 | WM170120-10 |
| ICBG1719 | Burkholderia | Atta sp. | 2341 | -22.84652, -48.43412 | Sao Paulo State | | 1/6/17 | LK170106-06 |
| ICBG1720 | Burkholderia | Atta sp. | 2341 | -22.84652, -48.43412 | Sao Paulo State | | 1/6/17 | LK170106-06 |
| ICBG1724 | Burkholderia | Atta sp. | 2340 | -22.84652, -48.43411 | Sao Paulo State | | 1/6/17 | LK170106-05 |
| ICBG849 | Burkholderia | Paratrachymyrmex cf. mandibularis | 2185 | S2° 34.818 W61° 02.045 | Anavilhanas, AM | | 1/23/17 | AR170123-03 |
| ICBG860 | Burkholderia | Paratrachymyrmex sp. | 2256 | S2° 16' 13.7 W61° 01' 05.3 | Anavilhanas, AM | | 1/24/17 | WM170124-08 |
| ICBG641 | Burkholderia | Mycetophylax cf. plaumanni | 2324 | S2° 55.808 W59° 58.479 | Ducke Reserve, AM | | 1/28/17 | TF170128-03 |
| ICBG652 | Burkholderia | Paratrachymyrmex | 2270 | n/a | Anavilhanas, AM | | 1/24/17 | CC170124-12 |
| ICBG663 | Burkholderia | Unidentified Attina | 2321 | S2° 55.808 W59° 58.479 | Ducke Reserve, AM | | 1/28/17 | WM170128-06 |
| ICBG664 | Burkholderia | Unidentified Attina | 2321 | S2° 55.808 W59° 58.479 | Ducke Reserve, AM | | 1/28/17 | WM170128-06 |
| ICBG665 | Burkholderia | Unidentified Attina | 2321 | S2° 55.808 W59° 58.479 | Ducke Reserve, AM | | 1/28/17 | WM170128-06 |
| ICBG666 | Burkholderia | Unidentified Attina | 2321 | S2° 55.808 W59° 58.479 | Ducke Reserve, AM | | 1/28/17 | WM170128-06 |
| ICBG667 | Burkholderia | Unidentified Attina | 2321 | S2° 55.808 W59° 58.479 | Ducke Reserve, AM | | 1/28/17 | WM170128-06 |
| ICBG668 | Burkholderia | Unidentified Attina | 2321 | S2° 55.808 W59° 58.479 | Ducke Reserve, AM | | 1/28/17 | WM170128-06 |
| ICBG847 | Burkholderia | Paratrachymyrmex | 2185 | S2° 34.818 W61° 02.045 | Anavilhanas, AM | | 1/23/17 | AR170123-03 |
| ICBG851 | Burkholderia | Apterostigma | 2149 | S2° 31' 23,4 W60° 49' 31,9 | Anavilhanas, AM | | 1/20/17 | WM170120-10 |
| ICBG859 | Burkholderia | Paratrachymyrmex | 2256 | S2° 16' 13.7 W61° 01' 05.3 | Anavilhanas, AM | | 1/24/17 | WM170124-08 |
| ICBG1733 | Burkholderia | Atta sp. | 2340 | -22.84652, -48.43411 | Sao Paulo State | | 1/6/17 | LK170106-05 |
| ICBG946 | Burkholderia | Apterostigma sp. | 2086 | S2° 32' 1.9 W60° 50' 6.1 | Anavilhanas, AM | | 1/19/17 | BP170119-02 |
| ICBG947 | Burkholderia | Apterostigma sp. | 2086 | S2° 32' 1.9 W60° 50' 6.1 | Anavilhanas, AM | | 1/19/17 | BP170119-02 |
| ICBG949 | Burkholderia | Apterostigma sp. | 2086 | S2° 32' 1.9 W60° 50' 6.1 | Anavilhanas, AM | | 1/19/17 | BP170119-02 |
| ICBG965 | Burkholderia | Apterostigma sp. | 2086 | S2° 32' 1.9 W60° 50' 6.1 | Anavilhanas, AM | | 1/19/17 | BP170119-02 |
| ICBG1008 | Burkholderia | Apterostigma | 2149 | S2° 31' 23,4 W60° 49' 31,9 | Anavilhanas, AM | | 1/20/17 | WM170120-10 |
| ICBG856 | Burkholderia | Apterostigma | 2149 | S2° 31' 23,4 W60° 49' 31,9 | Anavilhanas, AM | | 1/20/17 | WM170120-10 |
| ICBG961 | Burkholderia | Apterostigma | 2149 | S2° 31' 23,4 W60° 49' 31,9 | Anavilhanas, AM | | 1/20/17 | WM170120-10 |
| ICBG863 | Burkholderia | Paratrachymyrmex | 2256 | S2° 16' 13.7 W61° 01' 05.3 | Anavilhanas, AM | | 1/24/17 | WM170124-08 |
| ICBG864 | Burkholderia | Paratrachymyrmex | 2256 | S2° 16' 13.7 W61° 01' 05.3 | Anavilhanas, AM | | 1/24/17 | WM170124-08 |
| ICBG912 | Burkholderia | Apterostigma | 2311 | n/a | Ducke Reserve, AM | | 1/28/17 | CC170128-12 |
| ICBG1744 | Burkholderia | Atta sp. | 2340 | -22.84652, -48.43411 | Sao Paulo State | | 1/6/17 | LK170106-05 |
| SID20336 | Burkholderia | Atta cephalotes | 3962 | N10.43311, W84.01439 | La Selva Biological Station, CR | | 4/4/18 | CF180404-10 |
| SID20392 | Burkholderia | Paratrachymyrmex | 3963 | N10.43152, W84.02538 | La Selva Biological Station, CR | | 4/4/18 | CF180404-11 |
| SID20338 | Burkholderia | Paratrachymyrmex | 3963 | N10.43152, W84.02538 | La Selva Biological Station, CR | | 4/4/18 | CF180404-11 |
| SID20340 | Burkholderia | Atta cephalotes | 3964 | N10.43152, W84.02544 | La Selva Biological Station, CR | | 4/5/18 | CF180405-01 |
| SID20342 | Burkholderia | Atta cephalotes | 3964 | N10.43152, W84.02544 | La Selva Biological Station, CR | | 4/5/18 | CF180405-01 |
| SID20344 | Burkholderia | Atta cephalotes | 3965 | N10.42338, W84.00136 | La Selva Biological Station, CR | | 4/5/18 | CF180405-02 |
| SID20345 | Burkholderia | Atta cephalotes | 3965 | N10.42338, W84.00136 | La Selva Biological Station, CR | | 4/5/18 | CF180405-02 |
| SID20347 | Burkholderia | Atta cephalotes | 3967 | N10.42438, W84.00157 | La Selva Biological Station, CR | | 4/6/18 | CF180406-02 |
| SID20348 | Burkholderia | Atta cephalotes | 3967 | N10.42438, W84.00157 | La Selva Biological Station, CR | | 4/6/18 | CF180406-02 |
| SID20349 | Burkholderia | Atta cephalotes | 3967 | N10.42438, W84.00157 | La Selva Biological Station, CR | | 4/6/18 | CF180406-02 |
| SID20350 | Burkholderia | Atta cephalotes | 3967 | N10.42438, W84.00157 | La Selva Biological Station, CR | | 4/6/18 | CF180406-02 |
| SID20351 | Burkholderia | Atta cephalotes | 3967 | N10.42438, W84.00157 | La Selva Biological Station, CR | | 4/6/18 | CF180406-02 |
| SID20352 | Burkholderia | Atta cephalotes | 3967 | N10.42438, W84.00157 | La Selva Biological Station, CR | | 4/6/18 | CF180406-02 |
| SID20353 | Burkholderia | Atta cephalotes | 3967 | N10.42438, W84.00157 | La Selva Biological Station, CR | | 4/6/18 | CF180406-02 |
| SID20355 | Burkholderia | Sericomyrmex | 3969 | N10.43033, W084.0111 | La Selva Biological Station, CR | | 4/7/18 | CF180407-02 |
| SID20356 | Burkholderia | Sericomyrmex | 3969 | N10.43033, W084.0111 | La Selva Biological Station, CR | | 4/7/18 | CF180407-02 |
| SID20365 | Burkholderia | Atta cephalotes | 3974 | N10.43136, W084.00557 | La Selva Biological Station, CR | | 4/3/18 | HH180403-03 |
| SID20369 | Burkholderia | Paratrachymyrmex | 3985 | n/a | La Selva Biological Station, CR | | 4/3/18 | MP180403-05 |
| SID20372 | Burkholderia | Paratrachymyrmex | 3985 | n/a | La Selva Biological Station, CR | | 4/3/18 | MP180403-05 |
| SID20373 | Burkholderia | Paratrachymyrmex | 3985 | n/a | La Selva Biological Station, CR | | 4/3/18 | MP180403-05 |
| SID20374 | Burkholderia | Paratrachmyrmex | 3985 | n/a | La Selva Biological Station, CR | | 4/3/18 | MP180403-05 |
| SID20376 | Burkholderia | Mycetomoellerius | 3986 | N10.42960, W84.00996 | La Selva Biological Station, CR | | 4/4/18 | MP180404-01 |
| SID20377 | Burkholderia | Mycetomoellerius | 3986 | N10.42960, W84.00996 | La Selva Biological Station, CR | | 4/4/18 | MP180404-01 |
| SID20378 | Burkholderia | Mycetomoellerius | 3986 | N10.42960, W84.00996 | La Selva Biological Station, CR | | 4/4/18 | MP180404-01 |
| SID20379 | Burkholderia | Atta cephalotes | 3987 | N10.43044, W84.01105 | La Selva Biological Station, CR | | 4/4/18 | MP180404-02 |
| SID20382 | Burkholderia | Atta cephalotes | 3992 | N10.43193, W84.01485 | La Selva Biological Station, CR | | 4/4/18 | MP180404-07 |
| SID20383 | Burkholderia | Atta cephalotes | 3992 | N10.43193, W84.01485 | La Selva Biological Station, CR | | 4/4/18 | MP180404-07 |
| SID20384 | Burkholderia | Atta cephalotes | 3992 | N10.43193, W84.01485 | La Selva Biological Station, CR | | 4/4/18 | MP180404-07 |
| SID20385 | Burkholderia | Atta cephalotes | 3992 | N10.43193, W84.01485 | La Selva Biological Station, CR | | 4/4/18 | MP180404-07 |
| SID20386 | Burkholderia | Atta cephalotes | 3992 | N10.43193, W84.01485 | La Selva Biological Station, CR | | 4/4/18 | MP180404-07 |
| SID20387 | Burkholderia | Atta cephalotes | 3992 | N10.43193, W84.01485 | La Selva Biological Station, CR | | 4/4/18 | MP180404-07 |
| SID20389 | Burkholderia | Paratrachymyrmex | 3994 | N10.43152, W84.02544 | La Selva Biological Station, CR | | 4/5/18 | MP180405-02 |
| SID20390 | Burkholderia | Paratrachymyrmex | 3994 | N10.43152, W84.02544 | La Selva Biological Station, CR | | 4/5/18 | MP180405-02 |
| SID21199 | Burkholderia | Sericomyrmex | 4009 | N10.42531, W084.00162 | La Selva Biological Station, CR | | 3/31/19 | CF190331-05 |
| SID21200 | Burkholderia | Sericomyrmex | 4009 | N10.42531, W084.00162 | La Selva Biological Station, CR | | 3/31/19 | CF190331-05 |
| SID21201 | Burkholderia | Sericomyrmex | 4010 | N10.42471, W084.00176 | La Selva Biological Station, CR | | 3/31/19 | CF190331-07 |
| SID21202 | Burkholderia | Sericomyrmex | 4010 | N10.42471, W084.00176 | La Selva Biological Station, CR | | 3/31/19 | CF190331-07 |
| SID21203 | Burkholderia | Sericomyrmex | 4011 | N10.42955, W084.01048 | La Selva Biological Station, CR | | 4/1/19 | CF190401-01 |
| SID21204 | Burkholderia | Sericomyrmex | 4011 | N10.42955, W084.01048 | La Selva Biological Station, CR | | 4/1/19 | CF190401-01 |
| SID21205 | Burkholderia | Sericomyrmex | 4011 | N10.42955, W084.01048 | La Selva Biological Station, CR | | 4/1/19 | CF190401-01 |
| SID21206 | Burkholderia | Sericomyrmex | 4011 | N10.42955, W084.01048 | La Selva Biological Station, CR | | 4/1/19 | CF190401-01 |
| SID21207 | Burkholderia | Sericomyrmex | 4011 | N10.42955, W084.01048 | La Selva Biological Station, CR | | 4/1/19 | CF190401-01 |
| SID21208 | Burkholderia | Sericomyrmex | 4011 | N10.42955, W084.01048 | La Selva Biological Station, CR | | 4/1/19 | CF190401-01 |
| SID21209 | Burkholderia | Sericomyrmex | 4012 | N10.42955, W084.01048 | La Selva Biological Station, CR | adjacent to CF190401-01 | 4/1/19 | CF190401-02 |
| SID21210 | Burkholderia | Sericomyrmex | 4012 | N10.42955, W084.01048 | La Selva Biological Station, CR | | 4/1/19 | CF190401-02 |
| SID21211 | Burkholderia | Sericomyrmex | 4012 | N10.42955, W084.01048 | La Selva Biological Station, CR | | 4/1/19 | CF190401-02 |
| SID21212 | Burkholderia | Sericomyrmex | 4012 | N10.42955, W084.01048 | La Selva Biological Station, CR | | 4/1/19 | CF190401-02 |
| SID21213 | Burkholderia | Sericomyrmex | 4012 | N10.42955, W084.01048 | La Selva Biological Station, CR | | 4/1/19 | CF190401-02 |
| SID21214 | Burkholderia | Sericomyrmex | 4012 | N10.42955, W084.01048 | La Selva Biological Station, CR | | 4/1/19 | CF190401-02 |
| SID21215 | Burkholderia | Sericomyrmex | 4012 | N10.42955, W084.01048 | La Selva Biological Station, CR | | 4/1/19 | CF190401-02 |
| SID21216 | Burkholderia | Sericomyrmex | 4012 | N10.42955, W084.01048 | La Selva Biological Station, CR | | 4/1/19 | CF190401-02 |
| SID21217 | Burkholderia | Sericomyrmex | 4012 | N10.42955, W084.01048 | La Selva Biological Station, CR | | 4/1/19 | CF190401-02 |
