## Supplemental Table 2 for "*Burkholderia* from fungus gardens of fungus-growing ants produce antifungals that inhibit the specialized parasite *Escovopsis*"

| Organism (16S identification) | ID number | Ant Host | Est. size (bp) | %GC | CDS | contigs | N50 | average coverage | Species (ANIm percent identity) | JspeciesWS tetra correlation search top hit | BioProject | Whole Genome Accession | SRA Accession |
|---|---|---|---|---|---|---|---|---|---|---|---|---|---|
| Burkholderia | ICBG862 | Paratrachymyrmex | 7986442 | 67.96 | 7028 | 958 | 16317 | 30 | B. gladioli ATCC 10248 (.97) | B. gladioli (GCA_001527485) MSMB1756 (.9996) | PRJNA564151 | VWWG00000000 | SRR10136585 |
| Burkholderia | ICBG637 | Mycetophylax | 7912056 | 68.26 | 6903 | 848 | 17648 | 27 | B. gladioli ATCC 10248 (.97) | B. gladioli (GCA_001527485) MSMB1756 (.9997) | PRJNA564151 | VWWC00000000 | SRR10136598 |
| Burkholderia | ICBG647 | Mycetophylax | 8207449 | 68.13 | 7053 | 482 | 34978 | 51 | B. gladioli ATCC 10248 (.97) | B. gladioli (GCA_001527485) MSMB1756 (.9997) | PRJNA564151 | VWWD00000000 | SRR10136588 |
| Burkholderia | ICBG955 | Apterostigma | 7982062 | 68.29 | 6819 | 422 | 36080 | 38 | B. gladioli ATCC 10248 (.97) | B. gladioli (GCA_001527485) MSMB1756 (.9995) | PRJNA564151 | VWWH00000000 | SRR10136584 |
| Burkholderia | ICBG962 | Apterostigma | 7913770 | 67.74 | 7111 | 1225 | 12004 | 28 | B. gladioli ATCC 10248 (.97) | B. gladioli (GCA_001527485) MSMB1756 (.9991) | PRJNA564151 | VWWI00000000 | SRR10136583 |
| Burkholderia | ICBG1735 | Atta sp. | 8136338 | 66.59 | 7226 | 240 | 67746 | 39 | B. lata FL-7-5-30-S1-D0 (.98) | B. lata (GCA_001718575) FL-7-5-30-S1-D0 (.9997) | PRJNA564151 | VWWE00000000 | SRR10136587 |
| Burkholderia | ICBG1719 | Atta sp. | 8174181 | 66.41 | 7336 | 472 | 44435 | 49 | B. lata FL-7-5-30-S1-D0 (.98) | B. lata (GCA_001718575) FL-7-5-30-S1-D0 (.9979) | PRJNA564151 | VWWJ00000000 | SRR10136619 |
| Burkholderia | ICBG1720 | Atta sp. | 8137953 | 66.47 | 7287 | 347 | 52454 | 42 | B. lata FL-7-5-30-S1-D0 (.98) | B. lata (GCA_001718575) FL-7-5-30-S1-D0 (.9991) | PRJNA564151 | VWWK00000000 | SRR10136618 |
| Burkholderia | ICBG1724 | Atta sp. | 7497824 | 66.3 | 6781 | 423 | 40856 | 37 | B. ambifaria AMMD (.96) | B. ambifaria MEX-5 (.9977) | PRJNA564151 | VWWL00000000 | SRR10136617 |
| Burkholderia | ICBG849 | Paratrachymyrmex | 7232450 | 63.74 | 6330 | 75 | 212914 | 52 | B. sp. H160 (.89) | B. sp. H160 (.987) | PRJNA564151 | VWWF00000000 | SRR10136586 |
| Burkholderia | ICBG860 | Paratrachymyrmex | 7049452 | 63.7 | 6208 | 140 | 109899 | 41 | B. sp. H160 (.89) | B. sp. H160 (.9871) | PRJNA564151 | VWWN00000000 | SRR10136615 |
| Burkholderia | ICBG641 | Mycetophylax cf. plaumanni | 8127228 | 62.99 | 7066 | 78 | 264938 | 26 | Paraburkholderia eburnea (.89) | Paraburkholderia eburnea JCM 18070 (.9799) | PRJNA564151 | VWWM00000000 | SRR10136616 |
| Burkholderia | SID20336 | Atta cephalotes | 6956314 | 63.65 | 6128 | 279 | 50717 | 41 | B. sp. H160 (.89) | B. sp. H160 (.98612) | PRJNA603049 | SAMN13916280 | SRR10969351 |
| Burkholderia | SID20392 | Paratrachymyrmex | 8616253 | 65.53 | 8026 | 1029 | 16078 | 50 | B. lata 383 (.96) | B. contaminans MS14 (.98554) | PRJNA603049 | SAMN13916299 | SRR10969340 |
| Burkholderia | SID20340 | Atta cephalotes | 8302601 | 63.06 | 7268 | 89 | 186396 | 30 | P. eburnea JCM 18070 (.89) | Paraburkholderia eburnea JCM 18070 (.97875) | PRJNA603049 | SAMN13916282 | SRR10969339 |
| Burkholderia | SID20342 | Atta cephalotes | 7566269 | 63.46 | 6631 | 229 | 79636 | 42 | B. sp. H160 (.89) | B. sp. H160 (.98807) | PRJNA603049 | SAMN13916283 | SRR10969338 |
| Burkholderia | SID20344 | Atta cephalotes | 8385398 | 65.96 | 7630 | 379 | 48859 | 41 | B. lata 383 (.94) | B. contaminans MS14 (.99839) | PRJNA603049 | SAMN13916284 | SRR10969337 |
| Burkholderia | SID20345 | Atta cephalotes | 10369999 | 65.32 | 9527 | 537 | 50597 | 43 | B. lata 383 (.95) | B. cepacia RB-39 (.99733) | PRJNA603049 | SAMN13916285 | SRR10969336 |
| Burkholderia | SID20347 | Atta cephalotes | 8420472 | 64.34 | 7479 | 584 | 31192 | 27 | P. tropica Ppe8 (.96) | Paraburkholderia tropica Ppe8 (.99862) | PRJNA603049 | SAMN13916286 | SRR10969335 |
| Burkholderia | SID20349 | Atta cephalotes | 8258919 | 66.18 | 7543 | 375 | 49957 | 51 | B. cepacia RB-39 (.98) | B. cepacia RB-39 (.99959) | PRJNA603049 | SAMN13916287 | SRR10969334 |
| Burkholderia | SID20353 | Atta cephalotes | 9024411 | 65.01 | 8096 | 530 | 40572 | 24 | B. ubonensis MSMB0783 (.91) | B. sp. A9 (.99558) | PRJNA603049 | SAMN13916288 | SRR10969333 |
| Burkholderia | SID20355 | Sericomyrmex | 7680268 | 67.07 | 6926 | 374 | 45272 | 42 | B. seminalis FL-5-4-10-S1-D7 (.99) | B. cenocepacia 869T2 (.9979) | PRJNA603049 | SAMN13916289 | SRR10969332 |
| Burkholderia | SID20365 | Atta cephalotes | 7188778 | 65.9 | 6829 | 1629 | 7585 | 24 | B. pyrrocinia MSMB1755 (.94) | B. pyrrocinia MSMB1755 (.99069) | PRJNA603049 | SAMN13916291 | SRR10969348 |
| Burkholderia | SID20369 | Paratrachymyrmex | 10035004 | 61.64 | 8948 | 155 | 248827 | 34 | P. caribensis MBA4 (.89) | Paraburkholderia terrae NBRC 100964 (.99517) | PRJNA603049 | SAMN13916290 | SRR10969349 |
| Burkholderia | SID20373 | Paratrachymyrmex | 8286883 | 66.33 | 7483 | 402 | 46417 | 52 | B. lata FL-7-5-30-S1-D0 (.96) | B. lata FL-7-5-30-S1-D0 (.99909) | PRJNA603049 | SAMN13916292 | SRR10969347 |
| Burkholderia | SID20378 | Mycetomoellerius | 8570316 | 65.82 | 7884 | 1062 | 15127 | 12 | B. lata 383 (.96) | B. cepacia RB-39 | PRJNA603049 | SAMN13916293 | SRR10969346 |
| Burkholderia | SID20379 | Atta cephalotes | 7479548 | 67.34 | 6614 | 730 | 20798 | 43 | B. gladioli FDAARGOS 389 (.88) | B. glumae LMG 2196 =ATCC 33617 (.97387) | PRJNA603049 | SAMN13916294 | SRR10969345 |
| Burkholderia | SID20384 | Atta cephalotes | 9126732 | 65.8 | 8361 | 662 | 28217 | 17 | B. lata 383 (.98) | B. lata 383 | PRJNA603049 | SAMN13916296 | SRR10969343 |
| Burkholderia | SID20389 | Paratrachymyrmex | 9002784 | 62.56 | 8026 | 109 | 159274 | 58 | P. caribensis MBA4 (.89) | Paraburkholderia caribensis MBA4 (.99648) | PRJNA603049 | SAMN13916297 | SRR10969342 |
| Burkholderia | SID20390 | Paratrachymyrmex | 8491711 | 66.27 | 7651 | 413 | 42964 | 30 | B. lata FL-7-5-30-S1-D0 (.96) | B. lata FL-7-5-30-S1-D0 (.99831) | PRJNA603049 | SAMN13916298 | SRR10969341 |
