## Supplemental Table 3 for "*Burkholderia* from fungus gardens of fungus-growing ants produce antifungals that inhibit the specialized parasite *Escovopsis*"

| | n | mean | sd | ICBG1719 | ICBG1720 | ICBG1735 | ICBG1724 | SID20373 | SID20355 | SID20345 | SID20349 | SID20390 | SID20384 | SID20378 | SID20392 | SID20365 | ICBG637 | ICBG955 | ICBG849 | SID20336 | ICBG860 | SID20340 | SID21199 | SID21202 | SID21203 | SID21204 | SID21206 | SID21208 | SID21210 | SID21211 | SID21212 | SID21213 | SID21214 | SID21215 | SID21216 |
|---|---|---|---|---|---|---|---|---|---|---|---|---|---|---|---|---|---|---|---|---|---|---|---|---|---|---|---|---|---|---|---|---|---|---|---|
| ICBG1719 | 8 | 2.5079 | 0.15035 | | 0.8007 | 0.7993 | 0.3924 | 0.2717 | 0.0002 | <.0001 | <.0001 | <.0001 | <.0001 | <.0001 | <.0001 | <.0001 | <.0001 | <.0001 | <.0001 | <.0001 | <.0001 | <.0001 | <.0001 | <.0001 | <.0001 | <.0001 | <.0001 | <.0001 | <.0001 | <.0001 | <.0001 | <.0001 | <.0001 | <.0001 | <.0001 |
| ICBG1720 | 8 | 2.4541 | 0.15035 | 0.8007 | | 0.9986 | 0.2684 | 0.177 | 0.0005 | <.0001 | <.0001 | <.0001 | <.0001 | <.0001 | <.0001 | <.0001 | <.0001 | <.0001 | <.0001 | <.0001 | <.0001 | <.0001 | <.0001 | <.0001 | <.0001 | <.0001 | <.0001 | <.0001 | <.0001 | <.0001 | <.0001 | <.0001 | <.0001 | <.0001 | <.0001 |
| ICBG1735 | 8 | 2.4538 | 0.15035 | 0.7993 | 0.9986 | | 0.2676 | 0.1764 | 0.0005 | <.0001 | <.0001 | <.0001 | <.0001 | <.0001 | <.0001 | <.0001 | <.0001 | <.0001 | <.0001 | <.0001 | <.0001 | <.0001 | <.0001 | <.0001 | <.0001 | <.0001 | <.0001 | <.0001 | <.0001 | <.0001 | <.0001 | <.0001 | <.0001 | <.0001 | <.0001 |
| ICBG1724 | 8 | 2.6901 | 0.15035 | 0.3924 | 0.2684 | 0.2676 | | 0.8066 | <.0001 | <.0001 | <.0001 | <.0001 | <.0001 | <.0001 | <.0001 | <.0001 | <.0001 | <.0001 | <.0001 | <.0001 | <.0001 | <.0001 | <.0001 | <.0001 | <.0001 | <.0001 | <.0001 | <.0001 | <.0001 | <.0001 | <.0001 | <.0001 | <.0001 | <.0001 | <.0001 |
| SID20373 | 8 | 2.7423 | 0.15035 | 0.2717 | 0.177 | 0.1764 | 0.8066 | | <.0001 | <.0001 | <.0001 | <.0001 | <.0001 | <.0001 | <.0001 | <.0001 | <.0001 | <.0001 | <.0001 | <.0001 | <.0001 | <.0001 | <.0001 | <.0001 | <.0001 | <.0001 | <.0001 | <.0001 | <.0001 | <.0001 | <.0001 | <.0001 | <.0001 | <.0001 | <.0001 |
| SID20355 | 8 | 1.6995 | 0.15035 | 0.0002 | 0.0005 | 0.0005 | <.0001 | <.0001 | | <.0001 | <.0001 | 0.0002 | 0.0009 | <.0001 | <.0001 | 0.0022 | 0.0001 | 0.0021 | <.0001 | <.0001 | <.0001 | <.0001 | 0.0009 | 0.0853 | 0.1161 | 0.1341 | 0.0362 | <.0001 | 0.3252 | 0.1517 | <.0001 | 0.1861 | 0.2488 | 0.0015 | 0.118 |
| SID20345 | 8 | 0.8291 | 0.15035 | <.0001 | <.0001 | <.0001 | <.0001 | <.0001 | <.0001 | | 0.0002 | 0.7989 | 0.4755 | 0.6445 | 0.8248 | 0.3254 | 0.8856 | 0.3306 | 0.0042 | 0.0031 | 0.0003 | 0.0103 | 0.9759 | 0.0407 | 0.0282 | 0.0234 | 0.1402 | 0.7938 | 0.0099 | 0.0327 | 0.0207 | 0.0247 | 0.0091 | 0.5704 | 0.0276 |
| SID20349 | 8 | 0.0246 | 0.15035 | <.0001 | <.0001 | <.0001 | <.0001 | <.0001 | <.0001 | 0.0002 | | <.0001 | <.0001 | 0.0011 | 0.0005 | <.0001 | 0.0001 | <.0001 | 0.3761 | 0.4325 | 0.8893 | 0.2346 | 0.0025 | <.0001 | <.0001 | <.0001 | <.0001 | 0.0014 | <.0001 | <.0001 | 0.2434 | <.0001 | <.0001 | <.0001 | <.0001 |
| SID20390 | 8 | 0.8834 | 0.15035 | <.0001 | <.0001 | <.0001 | <.0001 | <.0001 | 0.0002 | 0.7989 | <.0001 | | 0.6462 | 0.4741 | 0.6341 | 0.4658 | 0.9116 | 0.4723 | 0.0019 | 0.0014 | 0.0001 | 0.0049 | 0.8117 | 0.0697 | 0.0497 | 0.0418 | 0.2101 | 0.619 | 0.0183 | 0.0554 | 0.011 | 0.0428 | 0.0175 | 0.7401 | 0.0488 |
| SID20384 | 8 | 0.9811 | 0.15035 | <.0001 | <.0001 | <.0001 | <.0001 | <.0001 | 0.0009 | 0.4755 | <.0001 | 0.6462 | | 0.2407 | 0.3502 | 0.7867 | 0.5688 | 0.7948 | 0.0004 | 0.0003 | <.0001 | 0.0011 | 0.54 | 0.1635 | 0.1229 | 0.1061 | 0.3941 | 0.3568 | 0.0495 | 0.1291 | 0.0031 | 0.1034 | 0.0501 | 0.9257 | 0.121 |
| SID20378 | 8 | 0.7309 | 0.15035 | <.0001 | <.0001 | <.0001 | <.0001 | <.0001 | <.0001 | 0.6445 | 0.0011 | 0.4741 | 0.2407 | | 0.8102 | 0.1492 | 0.5451 | 0.1522 | 0.0158 | 0.012 | 0.0017 | 0.0345 | 0.7289 | 0.0137 | 0.009 | 0.0073 | 0.0607 | 0.8683 | 0.003 | 0.0114 | 0.0582 | 0.0083 | 0.0025 | 0.3204 | 0.0088 |
| SID20392 | 8 | 0.782 | 0.15035 | <.0001 | <.0001 | <.0001 | <.0001 | <.0001 | <.0001 | 0.8248 | 0.0005 | 0.6341 | 0.3502 | 0.8102 | | 0.2287 | 0.715 | 0.2328 | 0.0081 | 0.0061 | 0.0008 | 0.0188 | 0.8803 | 0.0246 | 0.0166 | 0.0136 | 0.0954 | 0.9549 | 0.0057 | 0.02 | 0.0346 | 0.0149 | 0.005 | 0.4401 | 0.0162 |
| SID20365 | 8 | 1.0388 | 0.15035 | <.0001 | <.0001 | <.0001 | <.0001 | <.0001 | 0.0022 | 0.3254 | <.0001 | 0.4658 | 0.7867 | 0.1492 | 0.2287 | | 0.4009 | 0.9916 | 0.0001 | <.0001 | <.0001 | 0.0004 | 0.4046 | 0.2525 | 0.1956 | 0.1714 | 0.5383 | 0.2416 | 0.0836 | 0.1999 | 0.0014 | 0.1636 | 0.087 | 0.731 | 0.1929 |
| ICBG637 | 8 | 0.8598 | 0.15035 | <.0001 | <.0001 | <.0001 | <.0001 | <.0001 | 0.0001 | 0.8856 | 0.0001 | 0.9116 | 0.5688 | 0.5451 | 0.715 | 0.4009 | | 0.4068 | 0.0027 | 0.0019 | 0.0002 | 0.0068 | 0.8826 | 0.0554 | 0.039 | 0.0326 | 0.1771 | 0.6932 | 0.0141 | 0.0443 | 0.0145 | 0.0339 | 0.0133 | 0.664 | 0.0383 |
| ICBG955 | 8 | 1.0365 | 0.15035 | <.0001 | <.0001 | <.0001 | <.0001 | <.0001 | 0.0021 | 0.3306 | <.0001 | 0.4723 | 0.7948 | 0.1522 | 0.2328 | 0.9916 | 0.4068 | | 0.0001 | 0.0001 | <.0001 | 0.0005 | 0.4095 | 0.2485 | 0.1923 | 0.1684 | 0.5322 | 0.2456 | 0.082 | 0.1966 | 0.0014 | 0.1608 | 0.0852 | 0.7383 | 0.1896 |
| ICBG849 | 8 | 0.2133 | 0.15035 | <.0001 | <.0001 | <.0001 | <.0001 | <.0001 | <.0001 | 0.0042 | 0.3761 | 0.0019 | 0.0004 | 0.0158 | 0.0081 | 0.0001 | 0.0027 | 0.0001 | | 0.92 | 0.4555 | 0.7606 | 0.0206 | <.0001 | <.0001 | <.0001 | <.0001 | 0.0165 | <.0001 | <.0001 | 0.7277 | <.0001 | <.0001 | 0.0014 | <.0001 |
| SID20336 | 8 | 0.1919 | 0.15035 | <.0001 | <.0001 | <.0001 | <.0001 | <.0001 | <.0001 | 0.0031 | 0.4325 | 0.0014 | 0.0003 | 0.012 | 0.0061 | <.0001 | 0.0019 | 0.0001 | 0.92 | | 0.5182 | 0.6854 | 0.0166 | <.0001 | <.0001 | <.0001 | <.0001 | 0.0128 | <.0001 | <.0001 | 0.6591 | <.0001 | <.0001 | 0.001 | <.0001 |
| ICBG860 | 8 | 0.0543 | 0.15035 | <.0001 | <.0001 | <.0001 | <.0001 | <.0001 | <.0001 | 0.0003 | 0.8893 | 0.0001 | <.0001 | 0.0017 | 0.0008 | <.0001 | 0.0002 | <.0001 | 0.4555 | 0.5182 | | 0.2937 | 0.0036 | <.0001 | <.0001 | <.0001 | <.0001 | 0.0021 | <.0001 | <.0001 | 0.2992 | <.0001 | <.0001 | 0.0001 | <.0001 |
| SID20340 | 8 | 0.2781 | 0.15035 | <.0001 | <.0001 | <.0001 | <.0001 | <.0001 | <.0001 | 0.0103 | 0.2346 | 0.0049 | 0.0011 | 0.0345 | 0.0188 | 0.0004 | 0.0068 | 0.0005 | 0.7606 | 0.6854 | 0.2937 | | 0.0383 | <.0001 | <.0001 | <.0001 | 0.0002 | 0.0338 | <.0001 | <.0001 | 0.9473 | <.0001 | <.0001 | 0.0034 | <.0001 |
| SID21199 | 4 | 0.8213 | 0.21262 | <.0001 | <.0001 | <.0001 | <.0001 | <.0001 | 0.0009 | 0.9759 | 0.0025 | 0.8117 | 0.54 | 0.7289 | 0.8803 | 0.4046 | 0.8826 | 0.4095 | 0.0206 | 0.0166 | 0.0036 | 0.0383 | | 0.0813 | 0.0618 | 0.0537 | 0.1999 | 0.8492 | 0.0262 | 0.065 | 0.0559 | 0.0525 | 0.0268 | 0.6147 | 0.0609 |
| SID21202 | 6 | 1.3023 | 0.1736 | <.0001 | <.0001 | <.0001 | <.0001 | <.0001 | 0.0853 | 0.0407 | <.0001 | 0.0697 | 0.1635 | 0.0137 | 0.0246 | 0.2525 | 0.0554 | 0.2485 | <.0001 | <.0001 | <.0001 | <.0001 | 0.0813 | | 0.8879 | 0.8335 | 0.6581 | 0.0311 | 0.54 | 0.8515 | <.0001 | 0.7698 | 0.5928 | 0.1644 | 0.882 |
| SID21203 | 6 | 1.337 | 0.1736 | <.0001 | <.0001 | <.0001 | <.0001 | <.0001 | 0.1161 | 0.0282 | <.0001 | 0.0497 | 0.1229 | 0.009 | 0.0166 | 0.1956 | 0.039 | 0.1923 | <.0001 | <.0001 | <.0001 | <.0001 | 0.0618 | 0.8879 | | 0.9449 | 0.564 | 0.0218 | 0.6323 | 0.9579 | <.0001 | 0.8743 | 0.6937 | 0.126 | 0.994 |
| SID21204 | 6 | 1.354 | 0.1736 | <.0001 | <.0001 | <.0001 | <.0001 | <.0001 | 0.1341 | 0.0234 | <.0001 | 0.0418 | 0.1061 | 0.0073 | 0.0136 | 0.1714 | 0.0326 | 0.1684 | <.0001 | <.0001 | <.0001 | <.0001 | 0.0537 | 0.8335 | 0.9449 | | 0.5204 | 0.0182 | 0.6799 | 0.9895 | <.0001 | 0.9265 | 0.7454 | 0.1099 | 0.9508 |
| SID21206 | 5 | 1.1882 | 0.19017 | <.0001 | <.0001 | <.0001 | <.0001 | <.0001 | 0.0362 | 0.1402 | <.0001 | 0.2101 | 0.3941 | 0.0607 | 0.0954 | 0.5383 | 0.1771 | 0.5322 | <.0001 | <.0001 | <.0001 | 0.0002 | 0.1999 | 0.6581 | 0.564 | 0.5204 | | 0.1052 | 0.3128 | 0.5467 | 0.0006 | 0.4817 | 0.3413 | 0.3759 | 0.5592 |
| SID21208 | 6 | 0.769 | 0.1736 | <.0001 | <.0001 | <.0001 | <.0001 | <.0001 | <.0001 | 0.7938 | 0.0014 | 0.619 | 0.3568 | 0.8683 | 0.9549 | 0.2416 | 0.6932 | 0.2456 | 0.0165 | 0.0128 | 0.0021 | 0.0338 | 0.8492 | 0.0311 | 0.0218 | 0.0182 | 0.1052 | | 0.0079 | 0.025 | 0.0542 | 0.0191 | 0.0074 | 0.4383 | 0.0213 |
| SID21210 | 5 | 1.4604 | 0.19017 | <.0001 | <.0001 | <.0001 | <.0001 | <.0001 | 0.3252 | 0.0099 | <.0001 | 0.0183 | 0.0495 | 0.003 | 0.0057 | 0.0836 | 0.0141 | 0.082 | <.0001 | <.0001 | <.0001 | <.0001 | 0.0262 | 0.54 | 0.6323 | 0.6799 | 0.3128 | 0.0079 | | 0.6835 | <.0001 | 0.7591 | 0.9179 | 0.0533 | 0.6374 |
| SID21211 | 5 | 1.3506 | 0.19017 | <.0001 | <.0001 | <.0001 | <.0001 | <.0001 | 0.1517 | 0.0327 | <.0001 | 0.0554 | 0.1291 | 0.0114 | 0.02 | 0.1999 | 0.0443 | 0.1966 | <.0001 | <.0001 | <.0001 | <.0001 | 0.065 | 0.8515 | 0.9579 | 0.9895 | 0.5467 | 0.025 | 0.6835 | | <.0001 | 0.9195 | 0.7469 | 0.1306 | 0.9636 |
| SID21212 | 6 | 0.2933 | 0.1736 | <.0001 | <.0001 | <.0001 | <.0001 | <.0001 | <.0001 | 0.0207 | 0.2434 | 0.011 | 0.0031 | 0.0582 | 0.0346 | 0.0014 | 0.0145 | 0.0014 | 0.7277 | 0.6591 | 0.2992 | 0.9473 | 0.0559 | <.0001 | <.0001 | <.0001 | 0.0006 | 0.0542 | <.0001 | <.0001 | | <.0001 | <.0001 | 0.0073 | <.0001 |
| SID21213 | 5 | 1.3778 | 0.19017 | <.0001 | <.0001 | <.0001 | <.0001 | <.0001 | 0.1861 | 0.0247 | <.0001 | 0.0428 | 0.1034 | 0.0083 | 0.0149 | 0.1636 | 0.0339 | 0.1608 | <.0001 | <.0001 | <.0001 | <.0001 | 0.0525 | 0.7698 | 0.8743 | 0.9265 | 0.4817 | 0.0191 | 0.7591 | 0.9195 | <.0001 | | 0.828 | 0.106 | 0.8799 |
| SID21214 | 6 | 1.4338 | 0.1736 | <.0001 | <.0001 | <.0001 | <.0001 | <.0001 | 0.2488 | 0.0091 | <.0001 | 0.0175 | 0.0501 | 0.0025 | 0.005 | 0.087 | 0.0133 | 0.0852 | <.0001 | <.0001 | <.0001 | <.0001 | 0.0268 | 0.5928 | 0.6937 | 0.7454 | 0.3413 | 0.0074 | 0.9179 | 0.7469 | <.0001 | 0.828 | | 0.0549 | 0.6992 |
| SID21215 | 6 | 0.9597 | 0.1736 | <.0001 | <.0001 | <.0001 | <.0001 | <.0001 | 0.0015 | 0.5704 | <.0001 | 0.7401 | 0.9257 | 0.3204 | 0.4401 | 0.731 | 0.664 | 0.7383 | 0.0014 | 0.001 | 0.0001 | 0.0034 | 0.6147 | 0.1644 | 0.126 | 0.1099 | 0.3759 | 0.4383 | 0.0533 | 0.1306 | 0.0073 | 0.106 | 0.0549 | | 0.1241 |
| SID21216 | 6 | 1.3388 | 0.1736 | <.0001 | <.0001 | <.0001 | <.0001 | <.0001 | 0.118 | 0.0276 | <.0001 | 0.0488 | 0.121 | 0.0088 | 0.0162 | 0.1929 | 0.0383 | 0.1896 | <.0001 | <.0001 | <.0001 | <.0001 | 0.0609 | 0.882 | 0.994 | 0.9508 | 0.5592 | 0.0213 | 0.6374 | 0.9636 | <.0001 | 0.8799 | 0.6992 | 0.1241 | |
